# Biosynthesis of the Fungal Nonribosomal Peptide Penilumamide A and Biochemical Characterization of a Pterin-Specific Adenylation Domain

**DOI:** 10.1101/2022.08.30.505926

**Authors:** Stephanie C. Heard, Jaclyn M. Winter

## Abstract

We report the characterization of the penilumamide A biosynthetic gene cluster from the marine-derived fungus *Aspergillus flavipes* CNL-338. *In vitro* reconstitution studies demonstrated that three Plm nonribosomal peptide synthetases encoding four modules are required for constructing the lumazine-containing tripeptide. Further investigations using dissected adenylation domains determined substrate specificity for methionine and anthranilic acid and led to the first biochemical characterization of an adenylation domain with selectivity for a pterin-derived building block.

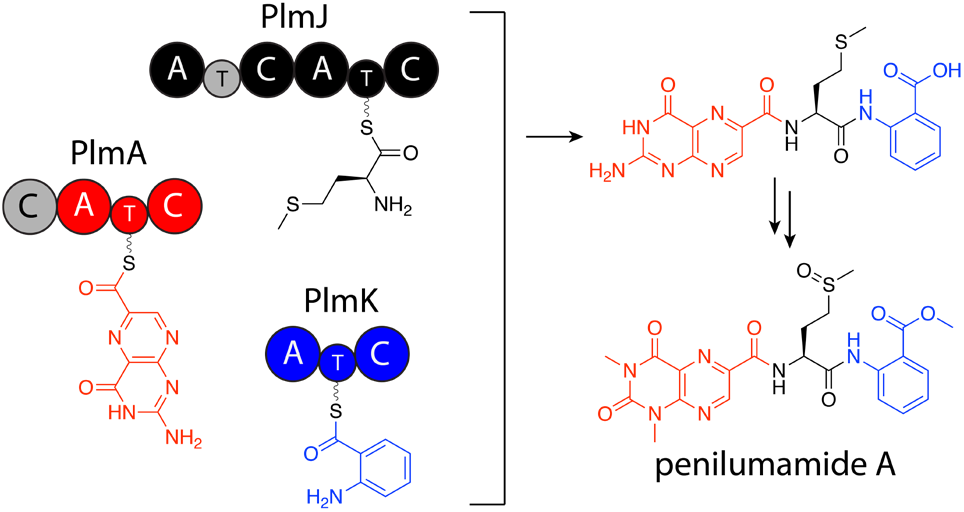

---

In the post-genomic era, natural product discovery has been enhanced through various genome mining and bioinformatic platforms. Strains that are likely to generate unique chemistry are prioritized due to our ability to identify putative biosynthetic clusters and predict building blocks and potential downstream modifications. Natural products produced by marine-derived microorganisms continue to be a treasure trove of novel bioactive small molecules^1–3^, including those of the well-established nonribosomal peptide class. Although the catalytic domains and mechanisms for peptide biosynthesis are similar between bacterial and fungal nonribosomal peptide synthetases (NRPSs), the ability to predict substrate specificity of building blocks in fungal systems is lacking compared to bacterial counterparts^4^. Thus, to enhance our ability to accurately predict nonribosomal peptides in fungal genomes, additional NPRS machinery needs to be biochemically characterized, especially those with selectivity for unprecedented nonproteinogenic building blocks.

In recent years, unique lumazine-containing nonribosomal peptides have been isolated from marine-derived *Aspergillus* strains (Figure 1). Penilumamide A (**1**), the first in its class, is a tripeptide containing a distinctive 1,3-dimethyl-lumazine-6-carboxylic acid functional group^5^, and this unique pterin-derived moiety is found in penilumamide analogs, usually exhibiting either 1-*N*-methylation or *N,N*-dimethylation. Additional structural differences with this suite of compounds includes variation in proteinogenic amino acid incorporation at the second position, as well as three different oxidation states of methionine, and addition of different aniline-derived C-terminal units such as anthranilic acid, methyl anthranilate, anthranilamide, or 2-aminophenyl isocyanide^6–11^. Despite the number of lumazine-containing peptides that have been isolated, there have been no biosynthetic investigations of the respective nonribosomal peptide machinery, and more importantly, no reports on adenylation domains with preference for lumazine- or pterin-derived building blocks. Herein, we report the biosynthetic pathway for **1** and the biochemical characterization of its NRPS machinery and corresponding adenylation domain substrate specificities. To the best of our knowledge, this is the first biochemical characterization of a fungal adenylation domain with selectivity for methionine and the first report of an adenylation domain that activates a pterin-derived building block.

**Figure 1.**
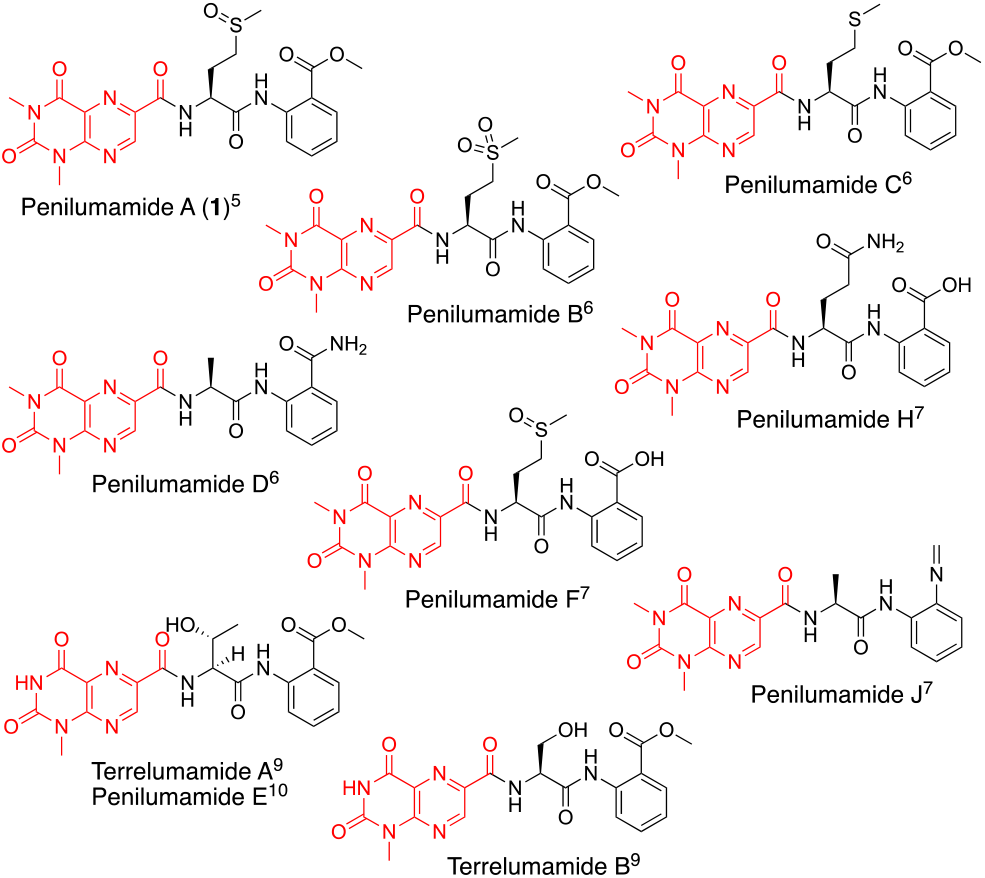
Structures of penilumamide A and other related lumazine-containing peptides. The lumazine-derived moieties are highlighted in red.

Nonribosomal peptides are a well-studied class of natural products that are typically assembled by large modular synthetases in an assembly-line like fashion. These megasynthetases provide a biosynthetic template where each module is typically responsible for the activation, incorporation and modification of proteinogenic or non-proteinogenic amino acid building blocks. Each module is made up of a minimal set of catalytic domains, namely adenylation, thiolation and condensation domains^12^. Adenylation (A) domains are responsible for selecting and activating building blocks, which then get loaded onto the phosphopantetheine moiety of a thiolation (T) domain. The tethered acyl substrate can then be delivered to the condensation (C) domain for extension with the upstream nascent peptide. To identify the biosynthetic machinery responsible for **1**, the 33 Mbp genome of *Aspergillus flavipes* CNL-338 was sequenced and assembled using SOAPdenovo2 and IDBA-UD software programs^13,14^. Initial automated annotation was carried out using antiSMASH^15^, which revealed 51 biosynthetic clusters, of which 23 are nonribosomal peptide-related. Additional genome mining using Blast+^16^ identified a 30 kb biosynthetic cluster, named the *plm* cluster, containing three NRPSs encoding four modules and eight genes dedicated to pterin biosynthesis (Figure 2A and Table S2). The eight non-NRPS genes within the *plm* biosynthetic cluster are hypothesized to convert guanosine triphosphate (GTP) to the final modified 1,3-dimethyllumazine-6-carboxylic acid unit, with the first two enzymatic steps predicted to be analogous to those in microbial folate biosynthesis.

**Figure 2.**
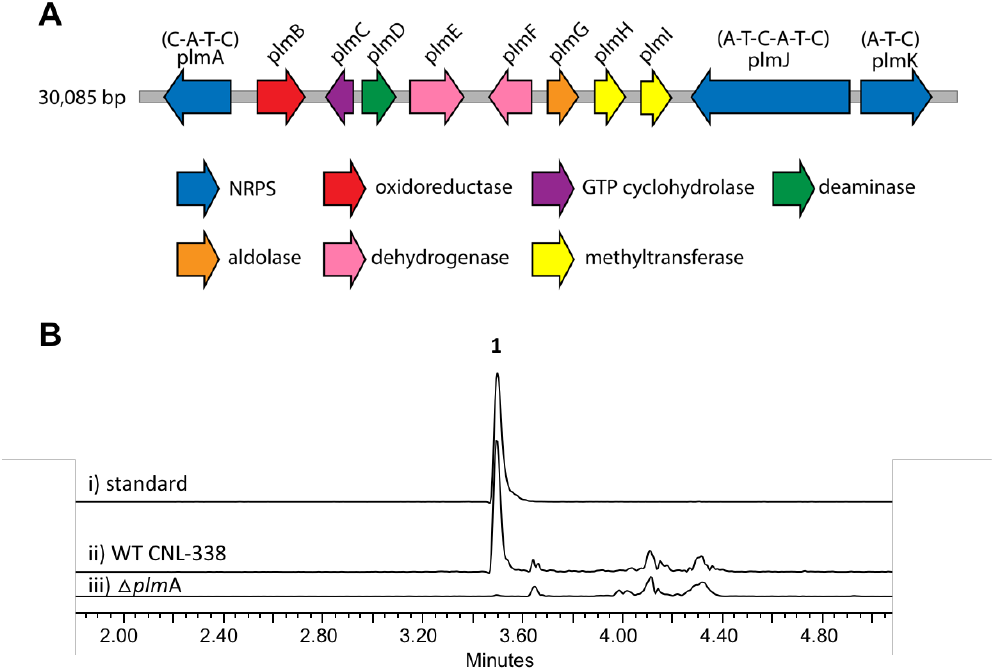
Organization and verification of the *plm* gene cluster in *A. flavipes* CNL-338. A) The *plm* cluster responsible for penilumamide A (**1**) production. The open reading frames are color-coded based on proposed function (Table S2). B) LC-MS analysis (EIC traces = 517 *m/z*) of i) a standard of **1** compared to crude extracts of ii) wildtype (WT) *A. flavipes* CNL-338 or iii) gene inactivation of the NRPS *plm*A.

Inactivation of *plm*A, a monomodular NRPS, confirmed the cluster’s role in synthesizing **1** (Figures 2B and S2)^17–19^. Interestingly, while penilumamide A is a linear tripeptide, closer inspection of the three NRPSs, PlmA, PlmJ and PlmK, revealed a total of four modules, indicative of a tetrapeptide product. Thus, the activities of the monomodular encoding PlmA and PlmK and dimodular encoding PlmJ were reconstituted *in vitro* to verify if all three NPRSs were required for assembly of **1**. PlmA, PlmJ and PlmK were solubly expressed as recombinant C-terminal hexahistidyl-tagged proteins from *Saccharomyces cerevisiae* BJ5464-NpgA^20,21^ and purified to near homogeneity in yields of 0.2 mg/L (Figure S3). Combinations of PlmA, PlmJ and PlmK were incubated with pterine-6-carboxylic acid, L-methionine and anthranilic acid before analysis by liquid chromatography-mass spectrometry (LC-MS). It should be noted that pterine-6-carboxylic acid was used as a commercially available alternative to the highly functionalized 1,3-dimethyllumazine-6-carboxylic acid building block produced by *A. flavipes* CNL-338, and no other tailoring enzymes were present in the *in vitro* assays. Thus, the expected product is the demethylated, pterin-containing penilumamide derivative (**2**) instead of **1**, and **2** was only observed when all three NRPSs, totaling four modules, were incubated together (Figure 3A).

**Figure 3.**
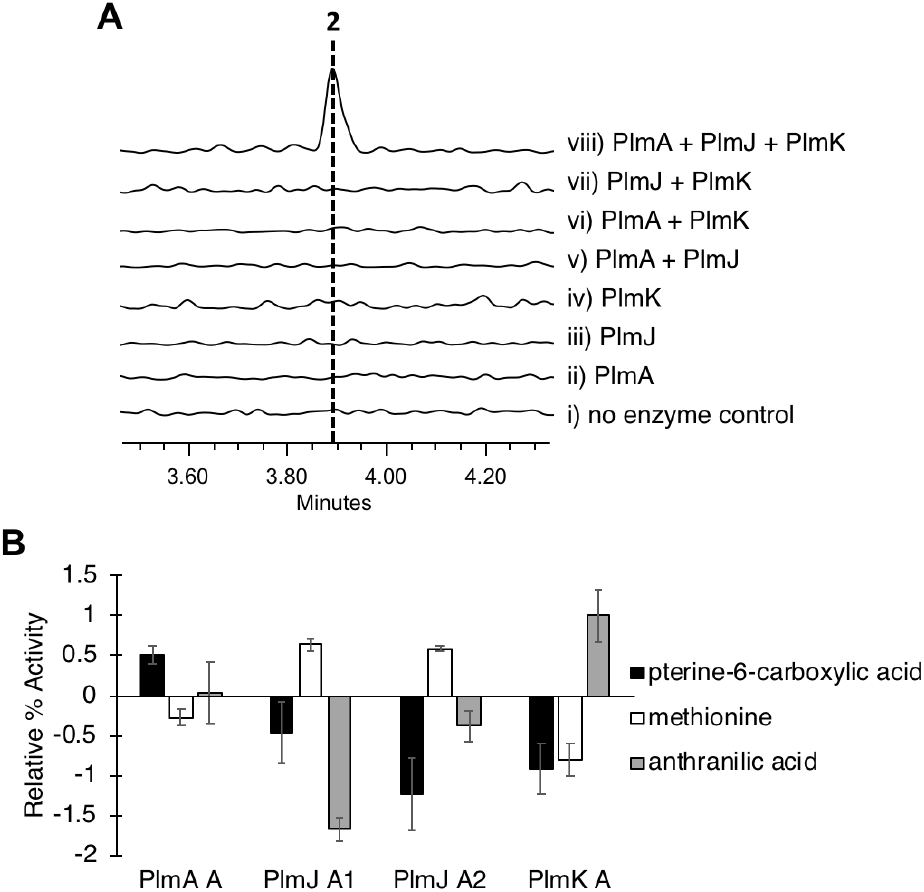
*In vitro* reconstitution of demethyl-pterin-penilumamide (**2**) and adenylation assays to determine substrate specificity of the four dissected adenylation domains found in PlmA, PlmJ and PlmK. A) LC-MS analysis (EIC = 458 *m/z*) of **2** when the pterine-6-carboxylic acid, L-methionine and anthranilic acid building blocks were (i) incubated together; (ii) incubated with PlmA; (iii) incubated with PlmJ; (iv) incubated with PlmK; (v) incubated with PlmA and PlmJ; (vi) incubated with PlmA and PlmK; (vii) incubated with PlmJ and PlmK; and (viii) incubated with PlmA, PlmJ and PlmK. B) Adenylation activity was determined through the malachite green/phosphate detection method^22^ using boiled enzyme controls, and all assays were run in triplicate. Relative % activity was measured as absorbance at 600 nm.

As the *in vitro* reconstitution assays showed that PlmA, PlmJ and PlmK were required for the production of **2**, detailed bioinformatic investigations of the signature motifs in each domain of the three Plm NRPSs was carried out to determine if any modules could be inactive (Figure S6). All four A domains in PlmA, PlmJ and PlmK were predicted to be active, and of the four Plm T domains, the first T domain in PlmJ (PlmJ_T1) is predicted to be inactive as it contains a glutamine residue instead of aspartate in the DxFFxLGGHSL motif. The first C domain in PlmJ however appears to be active as it contains the highly conserved HHxxxDG motif, whereas the N-terminal C domain in PlmA (PlmA_C1) is truncated and lacking key catalytic residues, and thus predicted to be inactive. As C domains have been posited as secondary gatekeepers to NRPS biosynthetic pathways^20^, downstream of the primary selectivity of A domains, a maximum-likelihood phylogenetic tree was constructed to aid in predicting the biosynthetic function of the Plm C domains, and perhaps module order (Table S5 and Figure S8). While the C domain in PlmK was shown to clade with terminal anthranilate-accepting domains, the second C domain in PlmJ (PlmJ_C2) clades with C domains that couple L-amino acid donors to anthranilate acceptors, thereby suggesting the order of NRPSs in the biosynthesis of **1** goes from PlmA to PlmJ to PlmK. To validate this order, and as all four A domains were predicted to be active, we investigated the substrate specificities of the individual A domains.

Adenylation domain sequences were excised from all three Plm encoding NRPSs based on predicted domain boundaries^21^ and expressed as N-terminal octahistidine-tagged proteins from *Escherichia coli* BL21 (DE3) (Figure S4). Using an established colorimetric phosphate detection method^22^, A domain activation assays were performed using pterine-6-carboxylic acid, anthranilic acid, formic acid, and the full panel of 20 proteinogenic amino acids (Figures 3B and S5). From these assays, the A domain in PlmA showed preference for pterine-6-carboxylic acid, whereas the second A domain in PlmJ (PlmJ_A2) activated methionine, and the A domain in PlmK was specific for anthranilic acid. Unexpectedly, the first A domain of PlmJ (PlmJ_A1) was also found to activate L-methionine to the same extent as PlmJ_A2, though only one methionine residue is present in **1**. However, these results support the necessary order of Plm NRPS modules predicted by C domain analysis.

As all four Plm A domains were biochemically characterized, the amino acid “specificity codes” of each domain were determined and compared to other fungal A domains to interrogate patterns denoting substrate selectivity (Table S3). It is well established in bacterial NRPSs that their substrate preferences can often be rationalized by 10 key amino acid residues that line the binding pocket of the A domain and act as a fingerprint for selectivity^23–25^. This work is the first report of an A domain, PlmA_A, activating a pterin-derived building block; its code DVMVLLMITK appears most similar to tryptophan-loading fungal A domains such as AnaPS A2 from acetylaszonalenin biosynthesis^26,27^. The A domain in PlmK, which was demonstrated *in vitro* to activate anthranilic acid, is comparable to anthranilate-activating fungal A domains^28^ such as PsyC A^29^. Both A domains of PlmJ were found to activate L-methionine *in vitro*, the first time this has been described in fungal NRPSs. Other methionine-incorporating A domains, such as NpsP8 from napsamycin biosynthesis^30^ and JahA A3 from jahnellamide biosynthesis^31^, have been predicted but lack biochemical confirmation. The specificity codes for PlmJ_A1 and PlmJ_A2 were determined to be SIVIVTAGTK and DVVLLLSSTK, respectively.

From detailed bioinformatic and biochemical investigations, the dimodular encoding PlmJ contributes only one methionine residue to penilumamide A, suggesting the first module may be catalytically inactive or skipped. In previous studies on fungal NRPS module skipping^32,33^, two possible mechanisms were suggested: either complete C-A-T module skipping via condensation by the downstream C domain, or chain transfer via the T domain of the skipped module. Based on our bionformatic analyses (Figure S6), the first of these mechanisms is more likely as the first T domain in PlmJ is the only T domain in the pathway that is lacking an initial aspartate residue in the conserved motif^34,35^, which may affect its ability to accept a peptide substrate despite the presence of the active site serine. Further, the A and C domains in the first module of PlmJ diverge from the other Plm domains in most bioinformatic analyses, and the predicted function of PlmJ_C1 as an anthranilate-accepting C domain does not correlate with the results of the PlmJ_A1 adenylation assays for methionine activation. All other modules in PlmA, PlmJ and PlmK have multiple lines of evidence supporting their proposed roles, making the first module of PlmJ incongruous in the pathway. Therefore, we predict that the entire A^1^-T^1^-C^1^ module of PlmJ is skipped in the biosynthesis of penilumamide A.

Based on bioinformatic data and *in vitro* biochemical assays, we propose that PlmA, which contains a truncated and inactive C domain at its N-terminus, activates and incorporates the unique pterin-derived building block, which originates from GTP via the GMC oxidoreductase *plm*B, the GTP cyclohydrolase I *plm*C, the aldehyde dehydrogenase *plm*E, the FAD-dependent oxidoreductase *plm*F, and the dihydroneopterin aldolase *plm*G. PlmJ is then able to extend the pterin-derived building block with L-methionine, and addition of anthranilic acid is catalyzed by PlmK (Figure 4). The tripeptide **2** is then offloaded from PlmK via hydrolysis, followed by deamination and methylation reactions facilitated by PlmD, and PlmH and PlmI, respectively, to afford **1**. However, we can’t rule out the possibility that deamination and *N*-methylation occur while the peptide is tethered to a thiolation domain, or if lumazine or 1,3-dimethyllumazine is the preferred building block of PlmA.

**Figure 4.**
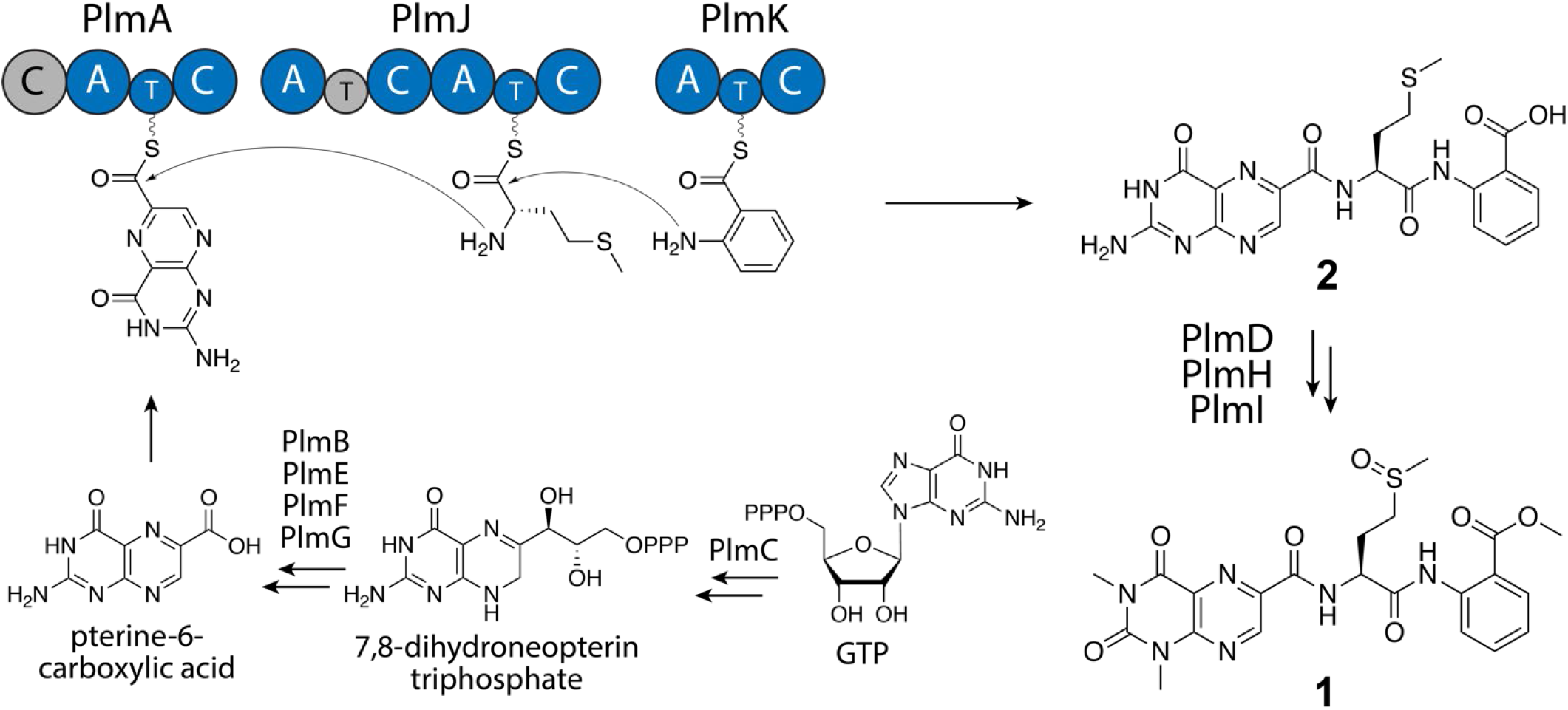
Proposed biosynthesis of penilumamide A (**1**). Biosynthesis of the pterin building block starting with GTP, and NPRS domain organization of PlmA, PlmJ and PlmK. NRPS domains consist of adenylation (A), thiolation (T) and condensation (C). Based on bioinformatic analysis, gray domains are proposed to be inactive.

In summary, we have identified and characterized the first biosynthetic cluster responsible for synthesizing a lumazine-containing natural product, penilumamide A, from the marine-derived fungus *A. flavipes* CNL-338. Using gene inactivation experiments and *in vitro* reconstitution assays, we have shown that all three *plm* NRPSs, encoding four modules, are required for the biosynthesis of the tripeptide, suggesting potential module skipping. Through detailed *in vitro* biochemical characterization assays, we determined the substrate specificity of the four *plm* NRPS adenylation domains, with bioinformatic analyses revealing the first “specificity codes” for methionine- and pterin-activating A domains. Altogether, this knowledge can be applied to other fungal NRPS systems and assist with bioinformatic-based predictions. Furthermore, penilumamides exhibit promising therapeutic potential as insulin sensitizing agents for the treatment of type II diabetes mellitus^9^. Understanding how subtle variations in chemical structure affect bioactivity will improve our bioengineering toolkit and expedite the development of penilumamides for use in patients.

## Supporting information

Penilumamide SI

## ASSOCIATED CONTENT

### Supporting Information

The Supporting Information is available free of charge on the ACS Publications website.

Experimental details, supplementary tables and figures, and phylogenetic analyses of adenylation and condensation domains (PDF).

## AUTHOR INFORMATION

### Author Contributions

S.C.H and J.M.W developed the project, carried out experiments, analyzed data, and wrote the manuscript. All authors have given approval to the final version of the manuscript.

### Notes

The authors declare there are no conflicts of interest. The penilumamide biosynthetic cluster is deposited under the GenBank accession number ON297683. The sequenced fungal internal transcribed spacer region for *Aspergillus flavipes* CNL-338 is deposited under accession number MT579592.

## ACKNOWLEDGMENT

We thank Professor William Fenical from Scripps Institution of Oceanography for providing *A. flavipes* CNL-338 and to Professor Yi Tang from UCLA for providing *S. cerevisiae* BJ5464 and the expression vector pXW55. S.C.H. thanks ARUP Laboratories and the Skaggs Foundation for graduate research fellowships. This work was supported by the Gordon and Betty Moore Foundation (GBMF7621, https://doi.org/10.37807/GBMF7621) and in part by the National Institutes of Health (1R01AI155694) to J.M.W.

