## Supplementary material for "Biosynthesis of the Fungal Nonribosomal Peptide Penilumamide A and Biochemical Characterization of a Pterin-Specific Adenylation Domain": Penilumamide SI

**Table of Contents**

**Experimental Methods	Page #**

Strains	S3

General molecular biology procedures	S3

Chemicals and spectroscopic procedures	S3

Fermentation conditions	S3

Isolation of fungal genomic DNA	S3

Genome sequencing, assembly and mining	S3

Construction of gene inactivation cassette for *plm*A	S4

Transformation and verification of disruption cassette in *A. flavipes* CNL-338	S4

LC-MS analysis of the Δ*plm*A mutant	S4

Cloning, expression and purification of *plm* NRPSs in yeast	S4

*In vitro* NRPS assay	S5

Cloning, expression and purification of *plm* A domains in bacteria	S6

*In vitro* A domain assays	S6

Bioinformatic analyses of *plm* enzyme domains	S7

**Supplementary Tables**

Table S1. Primers used in this work	S8

Table S2. Annotation of the *plm* gene cluster in *A. flavipes* CNL-338	S11

Table S3. Comparative analysis of adenylation domain residues that mediate amino acid specificity	S17

Table S4. Protein sequences used for 43 adenylation domain amino acid comparison	S19

Table S5. Protein sequences used for 53 condensation domain amino acid comparison	S22

Table S6. Protein sequences used for 200 condensation domain amino acid comparison	S25

**Supplementary Figures**

Figure S1. Verification of penilumamide A and derivative production	S10

Figure S2. Construction of inactivation cassette and verification of the Δ*plm*A mutant	S12

Figure S3. Plm NRPS reconstitution and expression in yeast	S13

Figure S4. SDS-PAGE of NRPS adenylation domains	S14

Figure S5. *In vitro* adenylation assays of the four NRPS modules	S15

Figure S6. Sequence alignments of *plm* thiolation, condensation and adenylation domains	S16

Figure S7. Phylogenetic analysis of NRPS adenylation domains	S21

Figure S8. Phylogenetic analysis of NRPS condensation domains	S24

Figure S9. Phylogenetic analysis of 200 NRPS condensation domains	S31

**Supplementary References**	S32

**Experimental Methods**

*Strains*

The fungal strain *Aspergillus flavipes* CNL-338 was obtained from Professor William Fenical at Scripps Institution of Oceanography. The strain was isolated as an endophyte from a red alga *Laurencia* sp. collected in the Bahamas^1^. Originally classified as a *Penicillium* sp., the strain identification was corrected upon morphological observation and fungal ITS sequence comparison (Accession number MT579592). *E. coli* BL21(DE3) was used to express the NRPS A domains from PlmA, PlmJ and PlmK. *Saccharomyces cerevisiae* strain BJ5464-NpgA (*MATα ura3-52 trp1 leu2-Δ1 his3Δ200 pep4::HIS3 prb1Δ1.6R can1 GAL*)^2^ was used to reconstitute and express the full NRPS enzymes PlmA, PlmJ and PlmK. This yeast strain contains the constitutive gene NpgA, a phosphopantetheine transferase from *Aspergillus nidulans,* integrated into the genome^3^, which allows for purification of *holo-*NRPS enzymes*.*

*General molecular biology procedures*

PCR reactions were carried out using AccuPrime Taq DNA polymerase (Invitrogen) and Phusion DNA polymerase (New England Biolabs). Restriction enzymes were purchased from New England Biolabs and used according to the manufacturer’s instructions. Primers were synthesized by Integrated DNA Technologies (Coralville, IA, USA). pCR®-blunt (Invitrogen) was used to construct recombinant DNA products, and DNA sequencing was performed by GeneWiz (South Plainfield, NJ, USA). DNA manipulation using standard techniques was performed in *E. coli* TOP10 (Invitrogen).

*Chemicals and spectroscopic procedures*

All solvents were purchased as HPLC grade or higher from Fisher Scientific (Waltham, MA, USA). Reverse phase LC-MS was performed using an Acquity Arc UHPLC/MS (Waters, Milford, MA, USA) in positive mode electrospray ionization with a Waters XBridge BEH C18 column (2.1 mm x 100 mm, 2.5 μm) fitted with the appropriate Waters VanGuard filter cartridge.

*Fermentation conditions*

For genomic DNA extraction, wild-type *A. flavipes* CNL-338 and the Δ*plm*A mutant were grown in 10 mL YPM media (0.2% yeast extract, 0.2% peptone, 0.4% mannitol) with 3.3% artificial sea salt (Instant Ocean, USA) in 10 x 35 mm petri dishes for 4 days at 30 °C under static conditions. For the Δ*plm*A mutant, 300 μg/mL of zeocin was added for antibiotic selection. For chemotype analysis between wild-type *A. flavipes* CNL-338 and the Δ*plm*A mutant, cultures were grown in 5 mL of YPM media with or without zeocin in 10 mL borosilicate glass tubes at 30 °C for 10 days at 170 rpm.

*Isolation of fungal genomic DNA*

Fungal cells were collected from the static culture into sterile 1.5 mL microcentrifuge tubes and lyophilized overnight. Cells were then broken into a fine powder by mechanical crushing. Lysis buffer (700 μL) (10 mM Tris-HCl pH 8.0, 20 mM EDTA, 0.5% SDS, 0.1 M LiCl) was added to each sample and inverted to afford a slurry, which was left at room temperature for five minutes. Phenol:chloroform:isoamyl alcohol (25:24:1) pH 8.0 (700 μL) was added to each sample, inverted to mix, and allowed to sit at room temperature for five minutes. Samples were spun to remove cell debris at 21,000 x g for 10 minutes at 4 °C. The aqueous top layer was transferred to a fresh microcentrifuge tube, where 500 μL of phenol:chloroform:isoamyl alcohol (25:24:1) pH 8.0 was added. Samples were centrifuged again, as described above, and the aqueous top layer was again transferred to a new microcentrifuge tube. Ethanol (1 mL of 95%) was added, tubes were inverted gently for five minutes at room temperature, and centrifuged again. The supernatant was aspirated and 400 μL of 70% ethanol was added to wash the pellet. After incubation at room temperature for five minutes, the samples were centrifuged at 21,000 x g for 2 minutes at room temperature. The supernatant was aspirated and pellets were dried at room temperature for ~15 minutes. Pellets were resuspended in 50 μL 10 mM Tris-HCl pH 8.0 and treated with RNase A (0.5 mg/mL) by incubating at 50 °C for 30 minutes. Genomic DNA (gDNA) was stored at 4 °C until needed.

*Genome sequencing, assembly and mining*

The genome of *A. flavipes* CNL-338 was sequenced at the High-Throughput Genomics Core at the Huntsman Cancer Institute at the University of Utah. A 180 bp PCR-free DNA library was constructed and sequenced using Illumina HiSeq (125 cycle paired-end) resulting in > 250 X coverage. Genome assembly was performed on the FutureSystems server using the SOAPdenovo (k-mer = 89) and IDBA-UD (k-mer = iterative) software packages^4,5^. Assembly of the 33 Mbp genome resulted in 1,899 contigs with an N_50_ value of 109,851 bp. Automated annotation was carried out using antiSMASH software^6^, which revealed 51 biosynthetic clusters, of which 23 are nonribosomal peptide-related. The assembled genome was also used to create a local BLAST database using Blast+ software (NCBI-BLAST®), and manual genome mining with the tblastn command confirmed the presence of a cluster containing three nonribosomal peptide synthatases encoding four modules as well as eight genes dedicated to pterin/lumazine biosynthesis. This resulting *plm* cluster was annotated and submitted to GenBank (Accession number ON2974468) (Figure 2A and Table S2).

*Construction of gene inactivation cassette for plmA*

To confirm the *plm* cluster’s involvement in penilumamide A biosynthesis, the NRPS *plm*A was targeted for gene inactivation using a disruption cassette (Figure S2). The zeocin resistance gene *Shble* and the constitutive tryptophan promoter PtrpC were designed by fusion PCR to disrupt *plm*A by antisense insertion^7^. The knockout cassette was constructed by attaching 2 kb of flanking homologous DNA located upstream and downstream of the targeted gene to the resistance marker. For construction of the *plm*A cassette, primer pairs plmA_KO_P1 and plmA_KO_P3 were used for the upstream region, while plmA_KO_P4 and plmA_KO_P6 were used for the downstream region (Table S1). The disruption cassette for inactivation of *plm*A was ligated into pCR®-blunt plasmid at 16 °C overnight and then transformed into *E. coli* TOP10. Its identity was confirmed by restriction enzyme digestion and DNA sequencing, and the cassette was amplified from pCR®-blunt using primers plmA_KO_P2 and plmA_KO_P5.

*Transformation and verification of disruption cassette in* A. flavipes *CNL-338*

Linearized inactivation cassette (10 µg) was transformed into *A. flavipes* CNL-338 as described previously^8,9^. Transformants were grown on stabilized minimal agar medium (1.2 M sorbitol, 1.5% agar, 1% dextrose, 5% nitrate salts, 0.1% trace elements) supplemented with 300 μg/mL zeocin. To confirm correct integration of the disruption cassette into the genome, gDNA was extracted and used as a template for PCR using primer pairs plmA_KO_P0 with PtrpC_R and zeocin_F with plmA_KO_P7 (Figure S2). These pieces both use an internal primer that anneals to either the promoter (PtrpC_R) or the resistance gene (zeocin_F) and an external primer that anneals either upstream or downstream of the region that was manipulated (Table S1). This effectively verifies not only the presence of the cassette in the mutant strain, but also the genomic location in the cluster. Wild-type (WT) *A. flavipes* CNL-338 gDNA was used as a control.

*LC-MS analysis of the* Δ*plm*A *mutant*

The Δ*plm*A *A. flavipes* CNL-338 mutant strain was cultured in 5 mL of YPM media with or without zeocin in 10 mL test tubes for 10 days at 170 rpm and extracted with two rounds of 1:1 volume ethyl acetate. Crude extracts were dried on a rotary evaporator, and the pellet was resuspended in 100 μL methanol. Extracts of the Δ*plm*A mutant were analyzed by LC-MS with a linear gradient of 5‒95% MeCN:H_2_O with 0.1% formic acid over 5 minutes followed by 95% MeCN for 1 minute at a flow rate of 0.6 mL/min. Mass data of the Δ*plm*A strain was compared by extracted ion chromatogram (EIC) to the wild-type *A. flavipes* CNL-338 as well as an authentic standard of penilumamide A (Figure 2B).

*Cloning, expression and purification of plm NRPSs in yeast*

The *plm*A sequence was predicted to contain two introns and thus was cloned in three overlapping pieces for intron-free reconstitution (4038 bp). Piece two (2154 bp) did not contain any introns and was amplified from gDNA using primers PlmA_F2 and PlmA_R2. Piece one (1116 bp) and piece three (1053 bp) were amplified from cDNA to remove predicted introns. RNA was extracted from a 3-day old YPM culture of WT *A. flavipes* CNL-338 using the RiboPure Yeast kit (Ambion). The manufacturer’s instructions were followed except contaminating gDNA was digested with DNase (2 U/μL) (Invitrogen) at 37 °C for 4 hours. cDNA was synthesized from total RNA for pieces one and three using SuperScript II Reverse Transcriptase (Invitrogen) with the reverse primers oligo_dT and PlmA_R3, respectively. The cDNA was used as a PCR template to amplify pieces one and three using primer pairs PlmA_F1 with PlmA_R1 and PlmA_F3 with PlmA_R3. All pieces were subcloned into pCR®-blunt (Invitrogen) for sequence verification. Piece two was re-amplified from the pCR-blunt construct with the same primer pairs as mentioned above, whereas pieces one and three were amplified using primer pairs PlmA_F1 with PlmA_yeast_R and PlmA_yeast_F with PlmA_R3, respectively, so that the amplified DNA contained the necessary overhangs for yeast recombination. A 2μ expression vector was linearized with PlmI and NdeI overnight at 37 °C. The three intron-free pieces of *plm*A were co-transformed with the linear vector into *S. cerevisiae* BJ5464-NpgA using the *S. c.* EasyComp™ Transformation Kit (Invitrogen) (Figure S3). The resulting expression plasmid pSHw_plmA, which places plmA under control of the ADH2 promoter, was sequence verified from transformants using primers pADH2, tADH2 and PlmA_ver1-PlmA_ver5.

A single confirmed transformant was inoculated into 3 mL SD_ct_ media (0.5% bacto cassamino acids technical grade, 2% dextrose) supplemented with adenine (0.02 mg/mL), tryptophan (0.02 mg/mL) and yeast nitrogen base solution (0.17% nitrogen base without amino acids, 0.5% ammonium sulfate) and grown for 3 days at 28 °C and 180 rpm. A 1 mL aliquot of this seed culture was used to inoculate 1 L of YPD media (1% yeast extract, 2% peptone) supplemented with 1% dextrose, and the culture was shaken at 28 °C and 180 rpm for 4 days. Yeast cells were harvested by centrifugation (3285 x g at 4 °C for 15 mins) and the pellet was resuspended in 30 mL lysis buffer (50 mM NaH_2_PO_4_, 150 mM NaCl, 10 mM imidazole, pH 8.0). Cells were sonicated on ice in one-minute intervals until homogenized. The lysate was cleared in two steps: first, the mixture was centrifuged at 37,156 x g and 4 °C for 1 hour, then the supernatant was passed through a 0.45 μm PVDF syringe filter. Ni-NTA agarose resin (2 mL) was added to the cleared lysate, and the solution was incubated on a rotator at 4 °C for 16 hours. Soluble PlmA (149.49 kDa) was purified by gravity-flow column chromatography using increasing concentrations of imidazole in buffer A (50 mM Tris-HCl, 500 mM NaCl, 20 mM-250 mM imidazole, pH 7.9). Purified protein was concentrated and buffer exchanged into buffer B (50 mM Tris-HCl, 2 mM EDTA, 2 mM DTT, 100 mM NaCl, pH 8.0) using an Amicon Ultracel 100,000 MWCO centrifugal filter (Merck Millipore Inc.) and stored in 10% glycerol at -80 °C until needed (Figure S3). The protein concentration was calculated to be 0.2 mg/L by Bradford assay using BSA as a standard.

The sequence for *plm*J was also predicted to contain two introns but due to its size (7128 bp), it was cloned in four overlapping pieces for intron-free reconstitution. Piece two (2934 bp) and piece three (2614 bp) did not contain any introns and were amplified from gDNA using primers pairs PlmJ_F2 with PlmJ_R2 and PlmJ_F3 with PlmJ_R3. Piece one (1149 bp) and piece four (1042 bp) were amplified from cDNA to remove predicted introns. RNA extraction and cDNA synthesis were performed as described above for *plm*A except reverse primers PlmJ_R1 and oligo_dT were used for pieces one and four, respectively. Pieces one and four were then amplified from the cDNA template using primer pairs PlmJ_F1 with PlmJ_R1 and PlmJ_F4 with PlmJ_R4, respectively. All pieces were subcloned into pCR®-blunt (Invitrogen) for sequence verification. Pieces two and three were re-amplified with the same primers from pCR-blunt, whereas pieces one and four were amplified to contain the necessary overhangs for recombination using primer pairs PlmJ_yeast_F with PlmJ_R1 and PlmJ_F4 with PlmJ_yeast_R, respectively. The co-transformation with linear expression vector was performed as described above for *plm*A. The resulting expression plasmid pSHw_plmJ was sequence verified from transformants using primers pADH2, tADH2 and PlmJ_ver1-PlmJ_ver10. Protein purification of PlmJ (261.74 kDa) was performed as described above for PlmA. The protein concentration was calculated to be 0.2 mg/L by Bradford assay using BSA as a standard.

The *plm*K sequence was predicted to contain one intron and thus was cloned in two overlapping pieces for intron-free reconstitution (3921 bp). Piece two (3121 bp) did not contain any introns and was amplified from gDNA using the primer pair PlmK_F2 with PlmK_R2. Piece one (1041 bp) was amplified from cDNA to remove the predicted intron. RNA extraction and cDNA synthesis were performed as described above for *plm*A except the reverse primer PlmK_R1 was used. All pieces were subcloned into pCR®-blunt (Invitrogen) for sequence verification. Both pieces of *plm*K were re-amplified with the same primers already containing the necessary overhangs for recombination. The co-transformation with linear expression vector was performed as described above for *plm*A. The resulting expression plasmid pSHw_plmK was sequence verified from transformants using primers pADH2, tADH2 and PlmK_ver1-PlmK_ver4. Protein purification of PlmK (144.97 kDa) was performed as described above for PlmA. The protein concentration was calculated to be 0.2 mg/L by Bradford assay using BSA as a standard.

*In vitro NRPS assay*

The NRPS machinery responsible for biosynthesizing penilumamide A was reconstituted *in vitro* using an established assay method from the Marahiel group^10^. In brief, each 100 μL reaction contained 50 mM Tris-HCl pH 8.0, 100 mM NaCl, 10 mM MgCl_2_, 2 mM ATP, 1 mM pterine-6-carboxylic acid, 1 mM l-methionine, 1 mM anthranilic acid, and 100 nM concentrations of PlmJ, PlmK and PlmA. The pterine-6-carboxylic acid substrate was used as a commercially available alternative to the highly functionalized 1,3-dimethyl-lumazine-6-carboxylic acid building block produced by *A. flavipes* CNL-338. PlmA, PlmJ and PlmK NRPSs were each incubated alone and in various combinations to determine the minimum number of modules required for tripeptide biosynthesis. Assays were incubated at 25 °C for 12 hours before extraction with a 1:1 volume of 99% ethyl acetate containing 1% acetic acid. The organic layer was resuspended in 50 μL methanol and analyzed by LC-MS with a linear gradient of 5‒95% MeCN:H_2_O with 0.1% formic acid over 5 minutes followed by 95% MeCN for 1 minute at a flow rate of 0.6 mL/min. Mass data was compared by extracted ion chromatogram (EIC) for the expected product across all assay conditions and compared to a no enzyme control reaction (Figure 3).

*Cloning, expression and purification of penilumamide A domains in bacteria*

*plm*A, *plm*J and *plm*K were analyzed by the Pfam 34.0 webtool^11^ to determine domain boundaries. All four adenylation (A) domains were not predicted to contain any introns. PlmA A and PlmJ A2 were amplified from gDNA of *A. flavipes* CNL-338. PlmA A (1485 bp) used the primer pair PlmA_A_F and PlmA_A_R, which contained EcoRI and NotI restriction sites, respectively. PlmJ A2 (1531 bp) used the primer pair PlmJ_A2_F and PlmJ_A2_R, which contained NcoI and NotI restriction sites, respectively. The two PCR products were subcloned into pCR®-blunt (Invitrogen) for sequence verification. The digested A domains were then ligated into the pHis8 expression vector^12^, which was prepared by digestion with the same corresponding restriction enzymes. The resulting expression plasmids pSHw_PlmA_A and pSHw_PlmJ_A2 were transformed into chemically competent *E. coli* TOP10 cells for sequence verification and *E. coli* BL21 (DE3) cells for protein expression. PlmJ A1 (1511 bp) and PlmK A (1524 bp) were codon optimized and synthesized by GeneWiz (South Plainfield, NJ, USA) into the pHis8 expression vector. The resulting expression plasmids pSHw_PlmJ_A1 and pSHw_PlmK_A, respectively, were transformed into chemically competent *E. coli* BL21 (DE3) cells for protein expression.

A single colony of each construct was used to inoculate 5 mL LB media (0.5% yeast extract, 1% peptone, 0.5% NaCl) supplemented with 50 μg/mL kanamycin and grown overnight at 37 °C and 170 rpm. These seed cultures were used to inoculate 1 L each of Terrific Broth (TB) (2.4% yeast extract, 1.2% tryptone, 0.004% glycerol) supplemented with 50 mg/L kanamycin. The cultures were incubated at 37 °C and 170 rpm until an OD_600_ of 1.0 was reached. Protein expression was induced with isopropyl-β-D-thiogalactopyranoside (IPTG) added to a final concentration of 0.1 mM, at which point the cultures were incubated at 16 °C for a further 16 hours. Bacteria cells were harvested by centrifugation (3285 xg at 4 °C for 15 mins) and the pellet was resuspended in 20 mL lysis buffer (50 mM Tris-HCl, 500 mM NaCl, 10 mM imidazole, pH 7.9). Cells were sonicated on ice in 30 second intervals until homogenized. The lysate was cleared by centrifugation at 32,914 x g and 4 °C for 30 minutes. Ni-NTA agarose resin (1 mL) was added to the supernatant, and the solution was incubated on a rotator at 4 °C for 16 hours. Soluble PlmA A (56 kDa), PlmJ A1 (58 kDa), PlmJ A2 (58 kDa) and PlmK A (57 kDa) were independently purified by gravity-flow column chromatography using increasing concentrations of imidazole in buffer A (50 mM Tris-HCl, 500 mM NaCl, 20 mM-250 mM imidazole, pH 7.9). Purified proteins were concentrated and buffer exchanged into buffer B (50 mM Tris-HCl, 2 mM DTT, pH 8.0) using Amicon Ultracel 50,000 MWCO centrifugal filters (Merck Millipore Inc.) and stored in 10% glycerol at -80 °C (Figure S3). The protein concentrations were calculated by Bradford assay using BSA as a standard (PlmA A = 17.3 mg/L, PlmJ A1 = 6.2 mg/L, PlmJ A2 = 12.8 mg/L and PlmK A = 11.5 mg/L).

*In vitro A domain assays*

The substrate loading of all four *plm* A domains was interrogated by an established colorimetric *in vitro* method from the Garneau-Tsodikova group^13^. This protocol uses the Malachite Green Phosphate Assay Kit (cat # POMG-25H) from BioAssay Systems (Hayward, CA, USA). In brief, all assays were performed in 96-well plates, and each 40 μL reaction contained 50 mM Tris-HCl pH 8.0, 100 mM NaCl, 15 mM MgCl_2_, 2.25 mM ATP, 0.2 U/mL inorganic pyrophosphatase, 3 mM of each substrate and 1 μM enzyme. The full panel of 20 proteinogenic amino acids were tested, along with formic acid, anthranilic acid and pterine-6-carboxylic acid. PlmA A, PlmJ A1, PlmJ A2 and PlmK A were independently incubated with all substrates at 25 °C for 1 hour before addition of 10 μL of the malachite green reagent. Color was allowed to develop for 15 minutes before absorbance was measured at 600 nm using a Molecular Devices SpectraMax M5 microplate reader. All assays were performed in triplicate. Negative controls included reactions where no amino acids were added for all enzymes and boiled enzyme reactions with all substrates (Figures 5 and S5).

*Bioinformatic analyses of* plm *enzyme domains*

PlmA, PlmJ and PlmK were analyzed with the Pfam 34.0 webtool^11^ to determine the boundaries of all corresponding adenylation (A), thiolation (T) and condensation (C) domains. Sequence alignments were made using the ClustalW algorithm and visualized using the ClustalX software to interrogate various motifs (Figure S6). An in depth bioinformatic comparison of all Plm domains was used in proposing the biosynthesis of penilumamide A (Figure 4). All phylogenetic trees were constructed and visualized using MEGA 6.0 software with the JTT model of amino acid substitution.

Pfam annotations indicated that across the four *plm* NRPS modules, there were five predicted C domains, which were added to an existing but modified dataset of canonical bacterial C domains^14,15^ along with several other unique fungal C domains. A maximum-likelihood phylogenetic tree was assembled containing a total of 200 C domain sequences (Figure S9 and Table S6). Due to a lack of clarity from the large bacterial C domain tree, a smaller and more relevant phylogeny was generated containing 53 C domain sequences, of which 34 are of fungal origin (Table S5 and Figure S8). It was also predicted that each of the four *plm* NRPS modules contained an A domain, which were aligned and compared to other bacterial and fungal adenylation domains. A maximum-likelihood phylogenetic tree was constructed containing 43 NRPS A domains, of which 32 are of fungal origin (Table S4 and Figure S7). PlmA_A clades closest to the alanine-loading domain TqaA_A3 from tryptoquialanine biosynthesis^16^ but is also closely related to three tryptophan-incorporating domains. PlmJ_A1 and PlmK_A both clade with a distinct group of anthranilate-activating enzymes, and PlmJ_A2 is most closely related to the phenylalanine-loading BenZ_A2 from benzomalvin biosynthesis^17^ and the valine-loading PsyA_A2 from psychrophilin biosynthesis^18^.

Further, in an effort to better understand substrate specificity in fungal A domains, all 43 sequences also had their 10 amino acid ”specificity codes” tabulated (Table S3)^19–21^. While not absolutely required, the first position of the code is usually aspartate, but it appears that at least in fungi, a significant portion (14 out of 43 sequences in Table S3) of A domains have substitutions to glycine, alanine, serine, threonine, and even proline.

**Table S1. Primers used in this work.**

| Primer name | Sequence (shown 5’-3’) |
| --- | --- |
| plmA KO P0 | cttcactacatgctacaggcc |
| plmA KO P1 | tctcctccaaacgcatgtctggtg |
| plmA KO P2 | cacgtacctcaatggacatgtggtcc |
| plmA KO P3 | ttcaatatcatcttctgtcgacggaaggatatttcaaggctcctgag |
| plmA KO P4 | gaaggctttaatttgcaagctgatgggctgaacatagccaaagcac |
| plmA KO P5 | gtcgcactgatcacatagcttgtagg |
| plmA KO P6 | cattcgaagcagcggtctagatgtgg |
| plmA KO P7 | tgccatgggtcagctt |
| PtrpC R | gtcgacagaagatgatattg |
| zeocin F | agcttgcaaattaaagcctt |
| oligo_dT | ttttttttttttttt |
| PlmA_F1 | catagcttccacatccacg |
| PlmA_R1 | ctaaccagttgaagattggg |
| PlmA_F2 | ccgtgaatatgtgtgttc |
| PlmA_R2 | gtagaggcatatcctgc |
| PlmA_F3 | atggctaccaagaaatcagc |
| PlmA_R3 | ctgaagatgtcctgccag |
| PlmA_yeast_F | tcaactatcaactattaactatatcgtaataccatatggctaccaagaaatcagc |
| PlmA_yeast_R | tgtcatttaaattagtgatggtgatggtgatgcacaccagttgaagattgggcca |
| PlmJ_F1 | atgggttcgaatgaatcccag |
| PlmJ_R1 | ggaatctggaattgactctgctg |
| PlmJ_F2 | tgccatcggcatccataa |
| PlmJ_R2 | cggtcataatcagtcagcatg |
| PlmJ_F3 | ctcgtggtaacatgtcaac |
| PlmJ_R3 | ctatgtcgttgagcagtacgg |
| PlmJ_F4 | actcccagatttcatcc |
| PlmJ_R4 | gatgaattctaccatg |
| PlmJ_yeast_F | tcaactatcaactattaactatatcgtaataccatatgggttcgaatgaatcccag |
| PlmJ_yeast_R | tgtcatttaaattagtgatggtgatggtgatgcacaactatattgccactgttc |
| PlmK_F1 | tcaactatcaactattaactatatcgtaataccatatgacgctttcagacttgtcg |
| PlmK_R1 | cctggcaatgtgaagataagatg |
| PlmK_F2 | ctggtacatttgtcccactctg |
| PlmK_R2 | tgtcatttaaattagtgatggtgatggtgatgcactgtatcttcatctcgaattgtcgca |
| pADH2_F | gcaaaacgtaggggcaaacaa |
| tADH2_R | gagctcggtaccctcga |
| PlmA_ver1 | gtatgatctcgtactatccgag |
| PlmA_ver2 | aagcataggaggagaactgc |
| PlmA_ver3 | gcctcaatgccaatcacacg |
| PlmA_ver4 | gattcaatcgacgcgatgaagg |
| PlmA_ver5 | ctagggttgccagtgtgattc |
| PlmJ_ver1 | ggggatacacactacaaacaag |
| PlmJ_ver2 | gatcctctcagagatgcaaacc |
| PlmJ_ver3 | catcggtaggcctacaaatg |
| PlmJ_ver4 | gcctttgaactgtgtgctcc |
| PlmJ_ver5 | ggctcagcaagagaagattg |
| PlmJ_ver6 | caaggaaacttgagcggc |
| PlmJ_ver7 | ttactctcgctggacagttg |
| PlmJ_ver8 | ctatttcgctaggtgacagcttg |
| PlmJ_ver9 | atccgaagctgtctacgttg |
| PlmJ_ver10 | gttcactgtctgaccgagaatgag |
| PlmK_ver1 | gcagtctcactctatcctatcg |
| PlmK_ver2 | cgaagtctactgcaccctga |
| PlmK_ver3 | cggtaggtgtcattgttgc |
| PlmK_ver4 | atggaccgaatggtgcgat |
| PlmA_A_F | **gaattc**cctgatgcaccggctgtttgtg |
| PlmA_A_R | **gcggccgc**ctagcggaggccaccccgatccacttt |
| PlmJ_A2_F | **ccatgg**caattgcagcaggtctggtcatc |
| PlmJ_A2_R | **gcggccgc**ctatcggaggaatttgcggtc |

EcoRI (GAATTC), NotI (GCGGCCG) and NcoI (CCATGG) restriction recognition sites are bolded and underlined

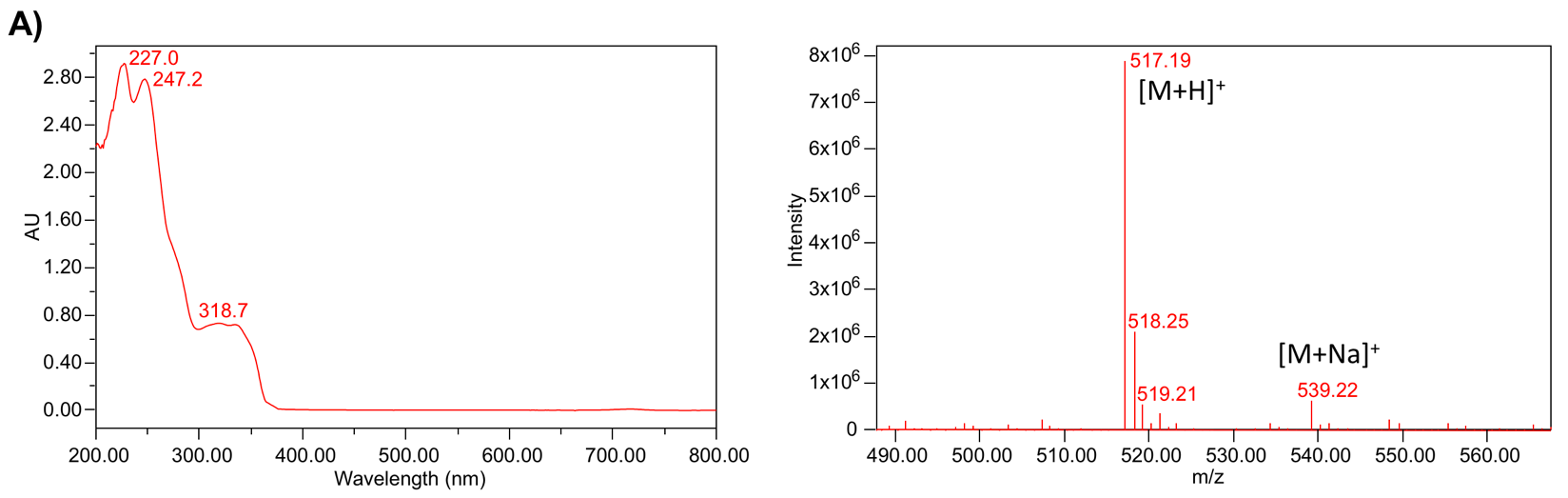

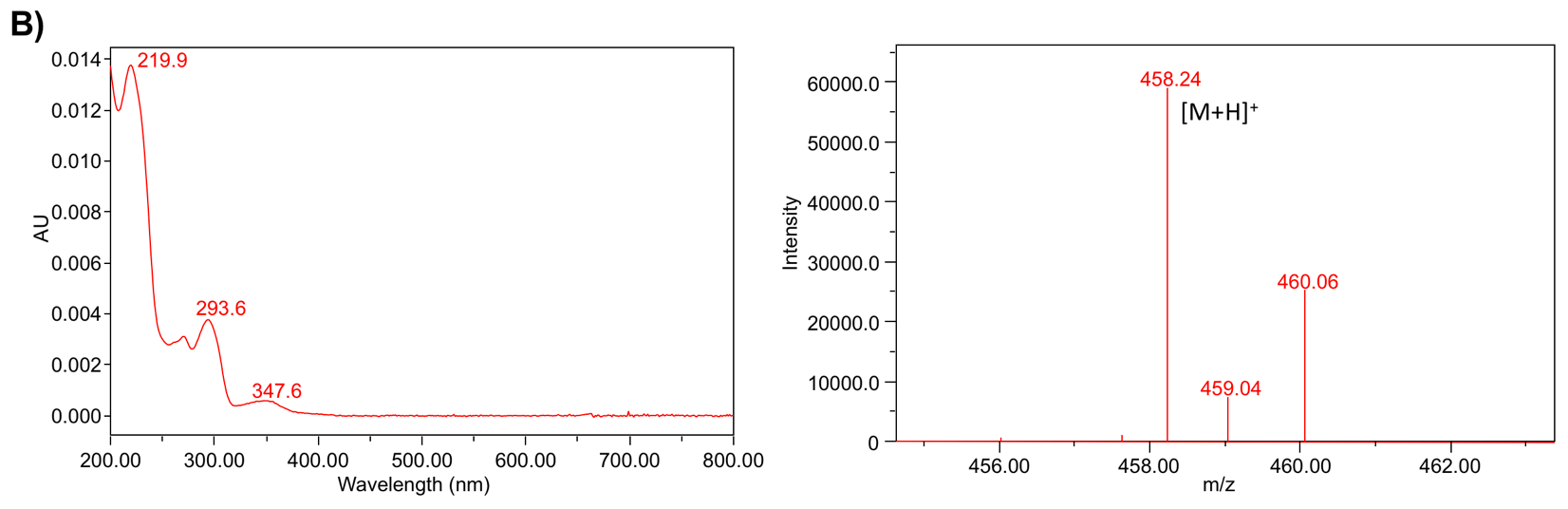

**Figure S1. Verification of penilumamide A from *A. flavipes* CNL-338 and the dimethyl-pterine-penilumamide from NRPS assays.** A) UV profile (left) and mass spectrum (right) of penilumamide A. B) UV profile (left) and mass spectrum (right) of the penilumamide derivative generated from NRPS assays in Figure 3A.

**Table S2. Annotation of the *plm* gene cluster in *Aspergillus flavipes* sp. CNL-338.** NRPS = nonribosomal peptide synthetase, C = condensation, A = adenylation, T = thiolation, GMC = glucose-methanol-choline, GTP = guanosine triphosphate, FAD = flavin adenine dinucleotide, SAM = S-adenosyl-l-methionine.

| **Gene**  **product** | **Total amino acids** | **Proposed**  **function** | **Sequence similarity (origin)** | **Identity/**  **similarity (%)** | **Accession**  **number** |
| --- | --- | --- | --- | --- | --- |
| PlmA | 1342 | NRPS (C_1_-A-T-C_2_) | *Aspergillus pseudoviridinutans* | 73/83 | GIJ86599.1 |
| PlmB | 591 | GMC oxidoreductase | *Penicillium* sp. *‘occitanis’* | 44/61 | PCG91499.1 |
| PlmC | 249 | GTP cyclohydrolase I | *Aspergillus pseudoviridinutans* | 80/86 | GIJ86601.1 |
| PlmD | 247 | Cytidine deaminase-like | *Aspergillus alliaceus* | 48/65 | XP_031896599.1 |
| PlmE | 487 | Aldehyde dehydrogenase | *Aspergillus avenaceus* | 51/67 | KAE8153438.1 |
| PlmF | 446 | FAD-dependent oxidoreductase | *Aspergillus flavus* AF70 | 45/64 | KOC15531.1 |
| PlmG | 313 | Dihydroneopterin aldolase/epimerase | *Penicillium roqueforti* FM164 | 51/72 | CDM28775.1 |
| PlmH | 244 | SAM-dependent methyltransferase | *Aspergillus flavus* | 40/57 | KAB8251571.1 |
| PlmI | 257 | SAM-dependent methyltransferase | *Aspergillus parasiticus* | 42/60 | KAB8205443.1 |
| PlmJ | 2375 | NRPS (A_1_-T_1_-C_1_-A_2_-T_2_-C_2_) | *Aspergillus udagawae* | 42/60 | GFF56995.1 |
| PlmK | 1313 | NRPS (A-T-C) | *Aspergillus udagawae* | 40/59 | GFF23642.1 |

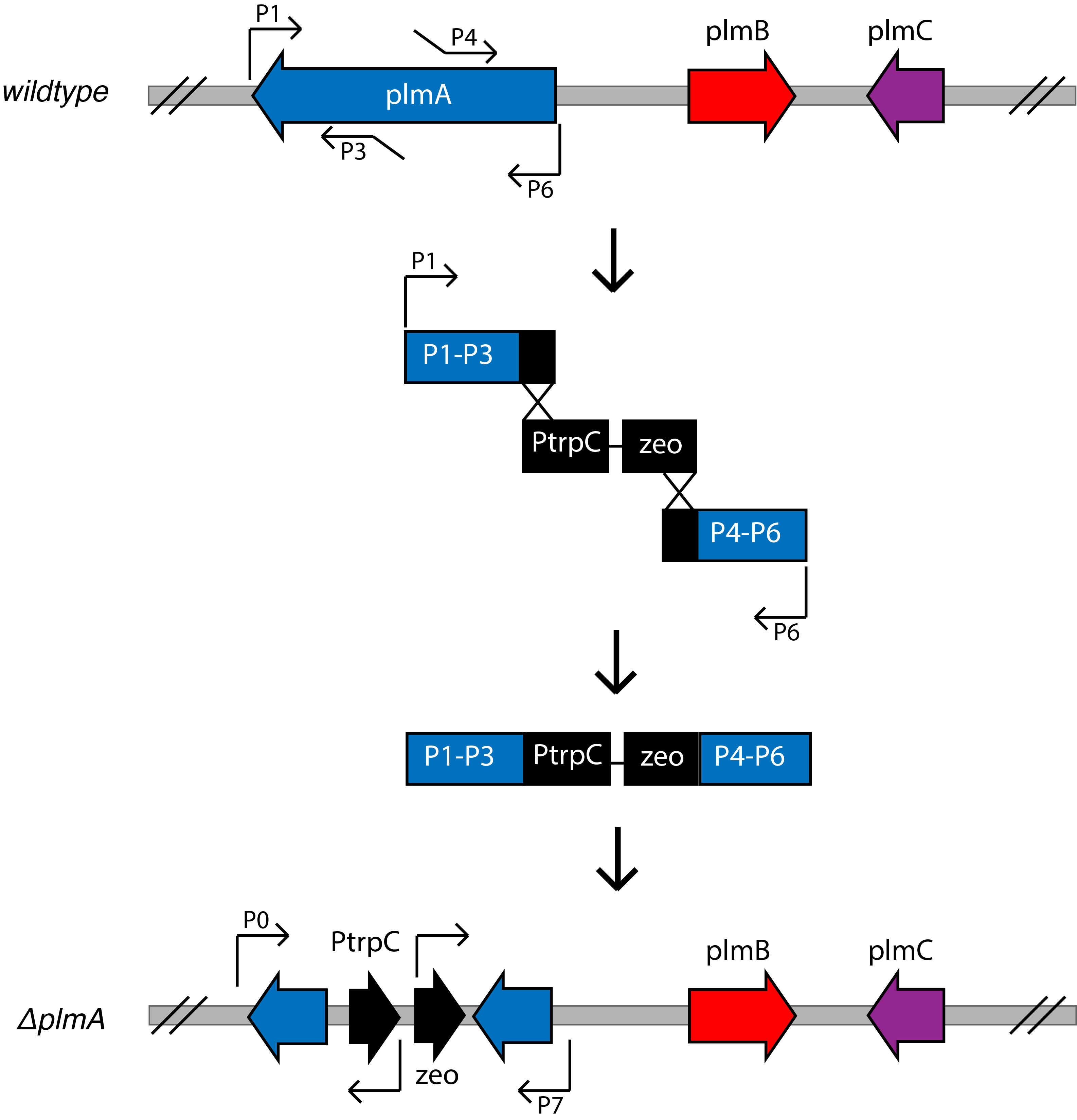

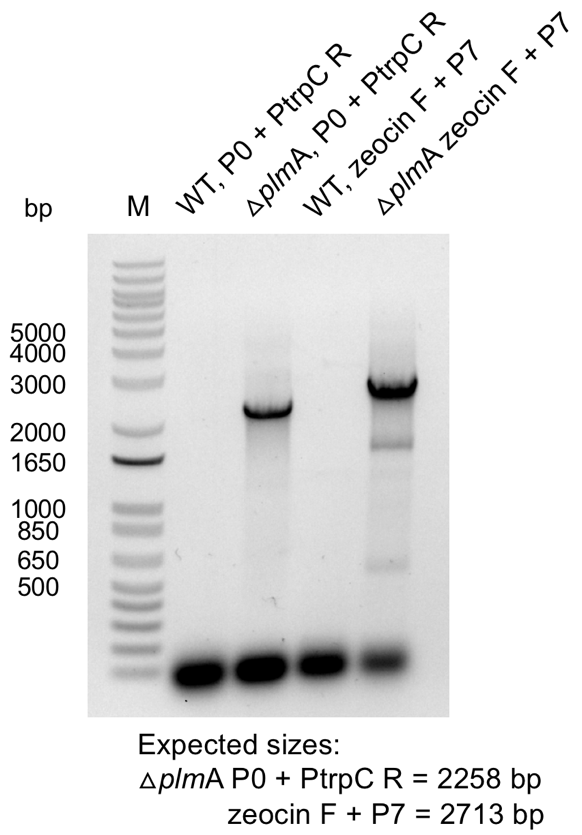

**Figure S2. Generation of the *plm*A NRPS gene inactivation cassette and integration into the genome by PCR verification.** Wild-type (WT) *A. flavipes* CNL-338 gDNA was used as a negative control.

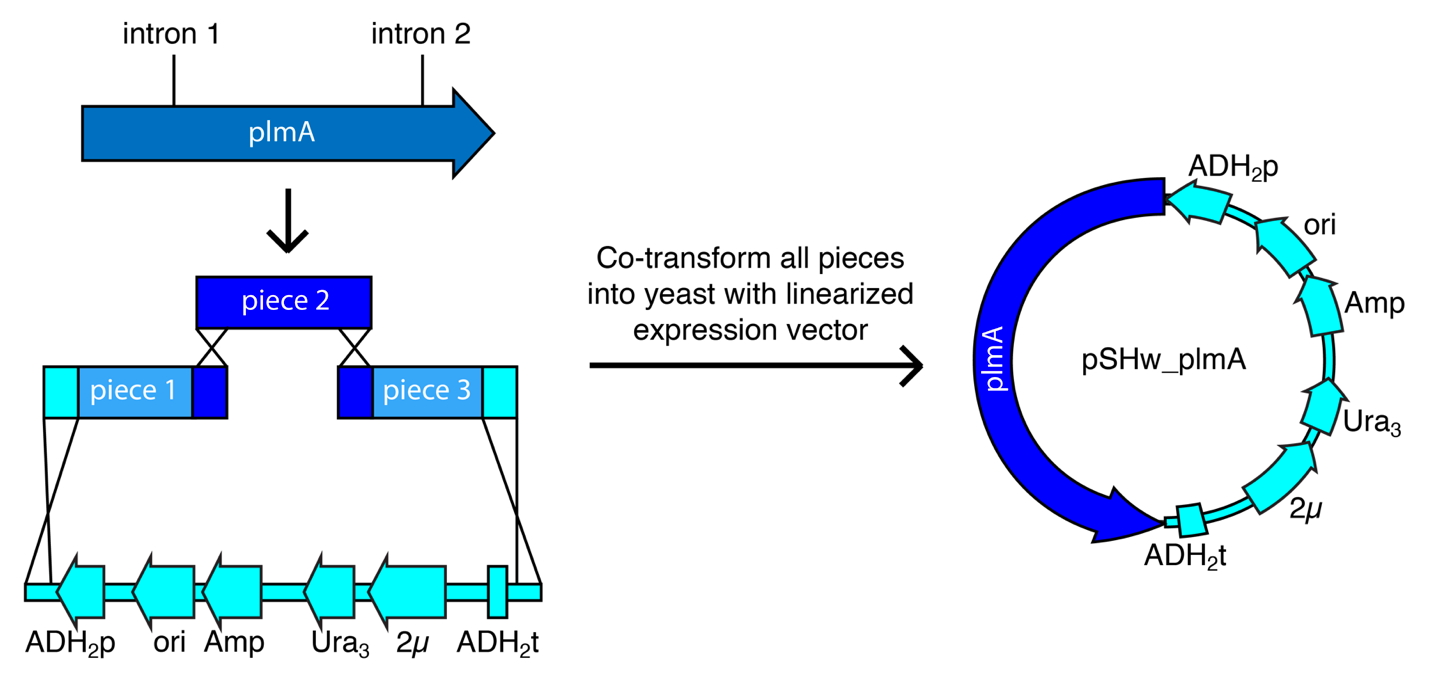

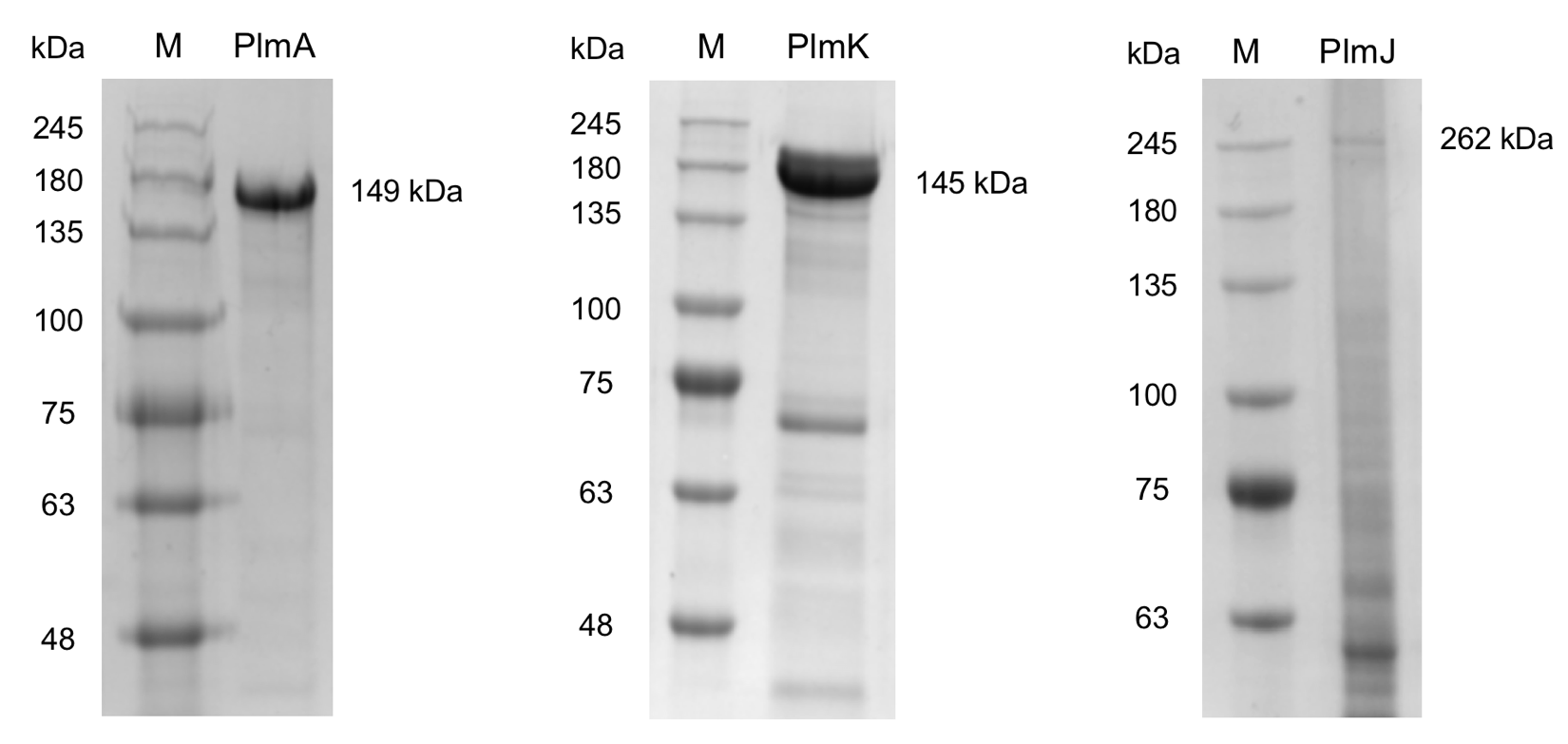

**Figure S3. Reconstituting PlmA, PlmJ and PlmK for expression in *S. cerevisiae BJ5464-NpgA*.** Overlapping regions between two neighboring DNA segments ranged from 140-187 bp and overlapping regions with the vector backbone were 35 bp. A representation for the reconstitution of intron-free PlmA is shown. PlmA, PlmJ and PlmK were expressed as C-terminal hexahistidyl-tagged proteins in *S. cerevisiae* BJ5464-NpgA and purified by Ni-NTA affinity chromatography to yield 0.2 mg/L, and analyzed for purity using an 8 % SDS-PAGE gel. Bluestain protein ladder 11-245 kDa (GoldBio) was used for all gels.

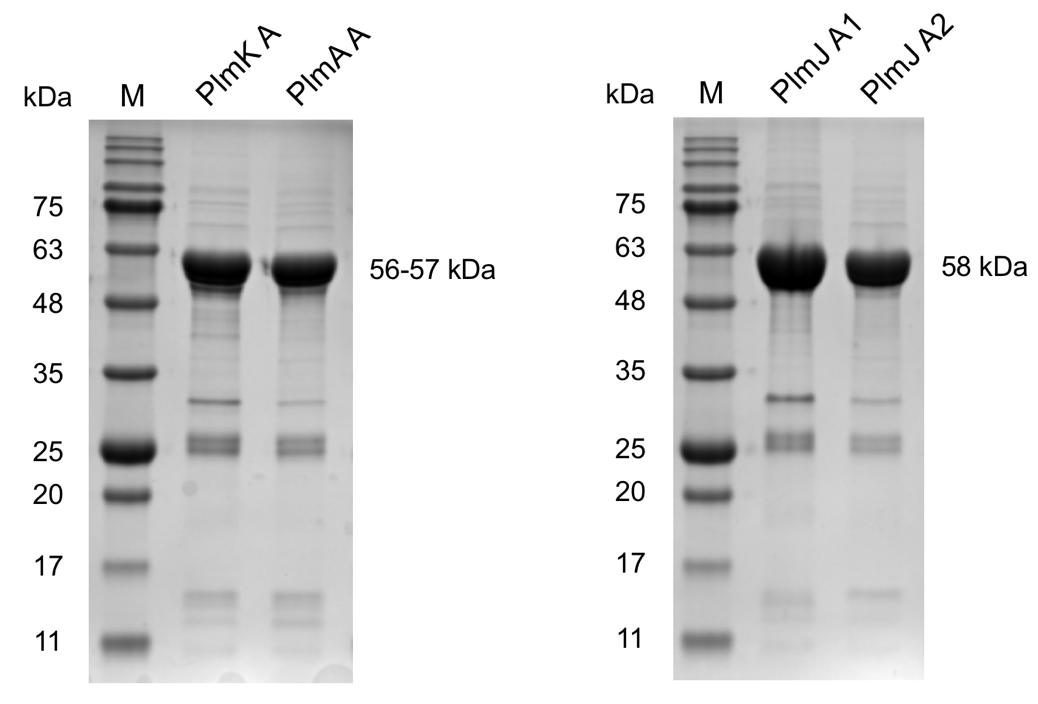

**Figure S4. SDS-PAGE of recombinant NRPS adenylation domains purified from *E. coli* BL21 (DE3).** Each dissected adenylation domain was expressed as a N-terminal octahistidyl-tagged protein, purified by Ni-NTA agarose affinity resin to yield between 6-17 mg/L, and analyzed for purity using a 12 % SDS-PAGE gel. Bluestain protein ladder 11-245 kDA (GoldBio) was used for all gels.

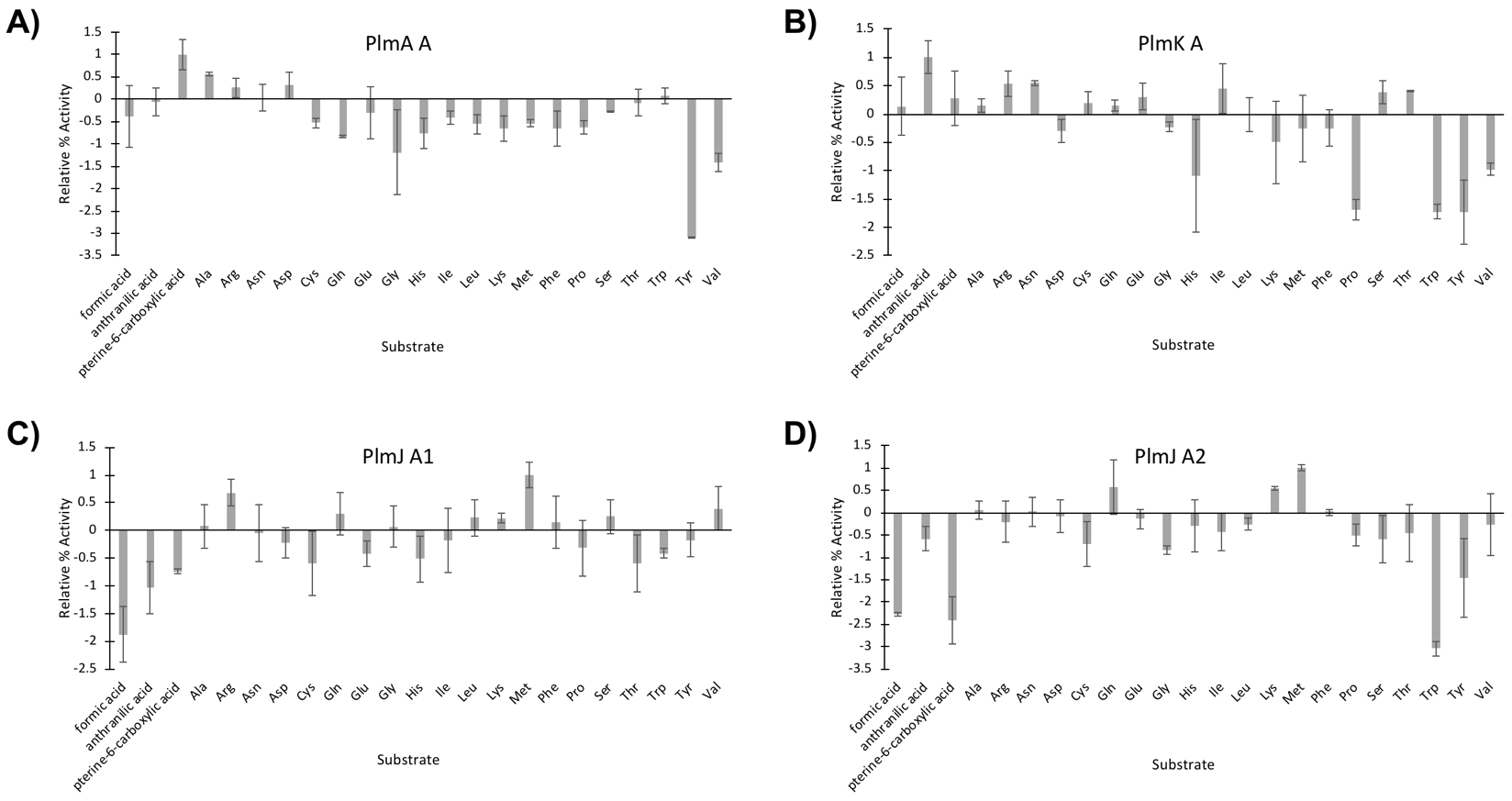

**Figure S5.** ***In vitro* adenylation assays to determine substrate loading of the four NRPS modules.** Data represents the mean of three replicates, and error bars represent the standard deviation. Adenylation activity was determined through the malachite green/phosphate detection method^13^ using a boiled enzyme control, and all assays were run in triplicate. Relative % activity was measured as absorbance at 600 nm.

**A)**

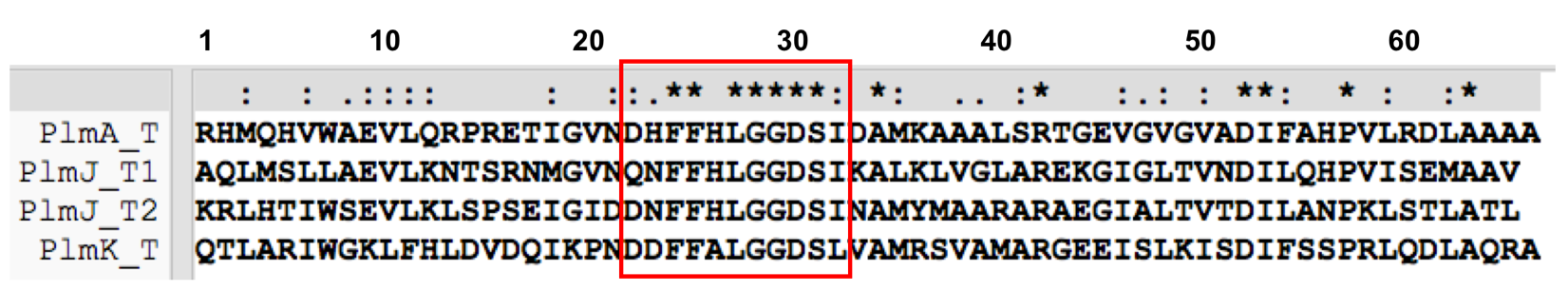

**B)**

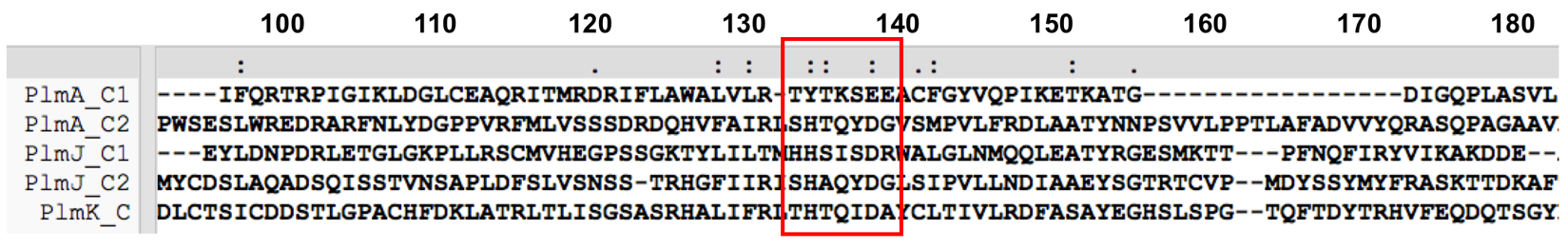

**C)**

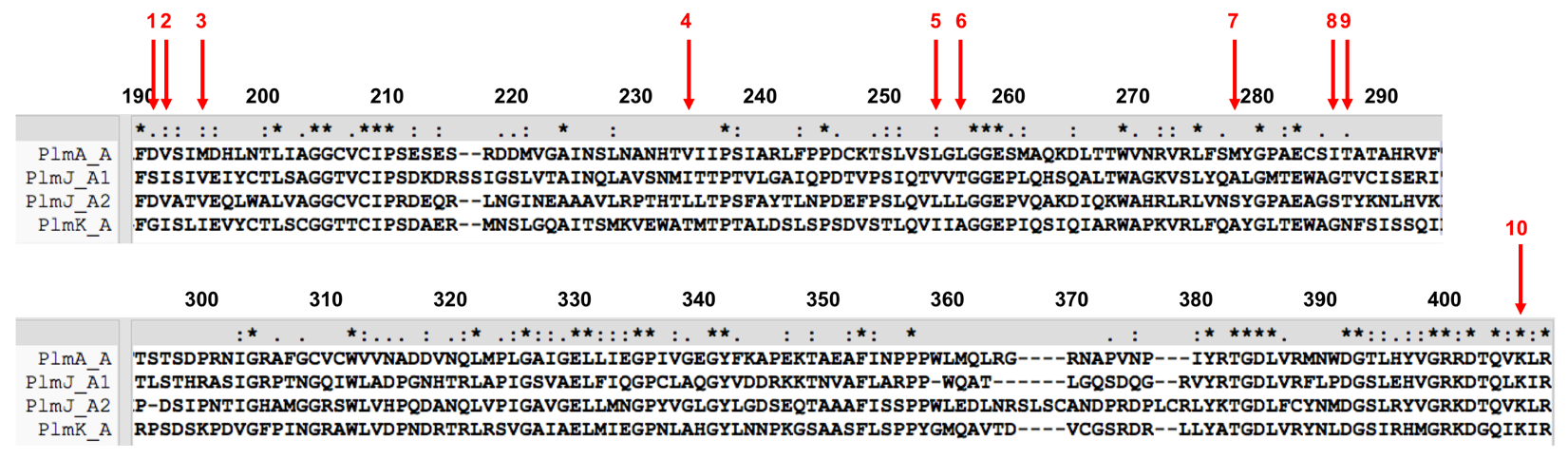

**Figure S6. Sequence alignments of thiolation, condensation and adenylation domains found in PlmA, PlmJ and PlmK.** A) Four thiolation (T) domains identified in the three *plm* NRPSs with the conserved DxFFxLGGHSL motif highlighted. B) Partial sequence of the five condensation (C) domains identified in the three *plm* NRPSs with the core HHxxxDG motif highlighted. The first C domain in PlmA (PlmA_C1) is predicted to be inactive as it does not contain any residues in this motif. C) Alignment of the four adenylation (A) domains with the 10 amino acid ”specificity code” denoted as 1-10. The amino acids are also listed in Table S3. The ClustalW algorithm was used for all alignments, and the ClustalX software was used for visualization.

**Table S3. Comparative analysis of adenylation domain residues that mediate amino acid specificity.** Residue positions are numbered according to AnaPS A2. Ant = anthranilic acid, Pip = pipecolate, pABA = para-aminobenzoic acid, l-Ky = l-kynurenine, AHBA = 3-amino-5-hydroxybenzoic acid, d-Hmp = d-2-hydroxy-3-methylpentanoic acid, AIA = 2-amioisobutyric acid.

| **Protein** | **Position 1**  **A.A. 190** | **Position 2**  **A.A. 191** | **Position 3**  **A.A. 194** | **Position 4**  **A.A. 231** | **Position 5**  **A.A. 251** | **Position 6**  **A.A. 253** | **Position 7**  **A.A. 275** | **Position 8**  **A.A. 283** | **Position 9 A.A. 284** | **Position 10**  **A.A. 392** | **Substrate** |
| --- | --- | --- | --- | --- | --- | --- | --- | --- | --- | --- | --- |
| **PlmA A** | **D** | **V** | **M** | **V** | **L** | **L** | **M** | **I** | **T** | **K** | **pterin** |
| **PlmJ A1** | **S** | **I** | **V** | **I** | **V** | **T** | **A** | **G** | **T** | **K** | **Met** |
| **PlmJ A2** | **D** | **V** | **V** | **L** | **L** | **L** | **S** | **S** | **T** | **K** | **Met** |
| **PlmK A** | **G** | **I** | **I** | **T** | **I** | **A** | **A** | **G** | **N** | **K** | **Ant** |
| **AnaPS A2** | **D** | **V** | **M** | **F** | **S** | **L** | **E** | **V** | **A** | **K** | **l-Trp** |
| **CheA A1** | **D** | **M** | **I** | **I** | **T** | **W** | **C** | **A** | **A** | **K** | **l-Trp** |
| **PsyA A1** | **G** | **A** | **T** | **F** | **L** | **L** | **G** | **S** | **A** | **K** | **l-Trp** |
| **TqaA A2** | **G** | **G** | **M** | **H** | **L** | **S** | **G** | **A** | **V** | **K** | **l-Trp** |
| **IvoA A** | **D** | **V** | **D** | **L** | **L** | **T** | **V** | **S** | **V** | **K** | **l-Trp** |
| **GrsA A1** | **D** | **A** | **W** | **T** | **I** | **A** | **A** | **I** | **C** | **K** | **l-Phe** |
| **CcsA A1** | **D** | **M** | **S** | **E** | **S** | **W** | **C** | **F** | **C** | **K** | **l-Phe** |
| **BenZ A2** | **G** | **M** | **N** | **V** | **L** | **L** | **G** | **G** | **V** | **K** | **l-Phe** |
| **Aba1 A3** | **D** | **A** | **W** | **V** | **L** | **S** | **G** | **I** | **Q** | **K** | **l-Phe** |
| **Aba1 A4** | **D** | **A** | **W** | **V** | **L** | **S** | **G** | **I** | **Q** | **K** | **l-Phe** |
| **GliP A1** | **D** | **G** | **G** | **I** | **I** | **L** | **A** | **T** | **C** | **K** | **l-Phe** |
| **AnaPS A1** | **G** | **A** | **L** | **F** | **L** | **I** | **A** | **G** | **V** | **K** | **Ant** |
| **AuaEII A** | **A** | **F** | **G** | **Y** | **C** | **S** | **G** | **H** | **I** | **K** | **Ant** |
| **BenZ A1** | **D** | **I** | **N** | **F** | **I** | **T** | **A** | **G** | **T** | **K** | **Ant** |
| **BenY A** | **D** | **M** | **F** | **I** | **V** | **T** | **L** | **G** | **M** | **K** | **Ant** |
| **PsyC A** | **D** | **I** | **I** | **L** | **I** | **S** | **A** | **G** | **I** | **K** | **Ant** |
| **TqaA A1** | **G** | **V** | **I** | **F** | **I** | **V** | **A** | **G** | **V** | **K** | **Ant** |
| **NanA A1** | **D** | **I** | **I** | **L** | **L** | **L** | **V** | **G** | **V** | **K** | **Ant** |
| **FkbP A** | **D** | **Y** | **Q** | **Y** | **L** | **Q** | **H** | **L** | **I** | **K** | **Pip** |
| **GetE A** | **D** | **V** | **Q** | **D** | **I** | **S** | **H** | **M** | **V** | **K** | **Pip** |
| **RapP A** | **D** | **Y** | **Q** | **Y** | **L** | **Q** | **H** | **L** | **V** | **K** | **Pip** |
| **SnbA A** | **P** | **F** | **P** | **S** | **L** | **V** | **V** | **L** | **T** | **K** | **Pip** |
| **SnbDE A3** | **D** | **F** | **Q** | **F** | **I** | **Q** | **V** | **A** | **V** | **K** | **Pip** |
| **PsyA A2** | **D** | **M** | **V** | **F** | **L** | **L** | **L** | **G** | **I** | **K** | **l-Val** |
| **Aba1 A2** | **G** | **A** | **W** | **M** | **L** | **A** | **A** | **I** | **L** | **K** | **l-Val** |
| **Aba1 A7** | **D** | **A** | **W** | **M** | **L** | **A** | **A** | **I** | **L** | **K** | **l-Val** |
| **Aba1 A9** | **D** | **A** | **W** | **M** | **L** | **A** | **A** | **I** | **L** | **K** | **l-Val** |
| **NpsP8** | **T** | **G** | **L** | **I** | **V** | **V** | **I** | **C** | **V** | **K** | **l-Met** |
| **Aba1 A5** | **D** | **V** | **W** | **V** | **M** | **S** | **A** | **I** | **Q** | **K** | **l-Pro** |
| **Aba1 A6** | **D** | **A** | **L** | **V** | **L** | **I** | **V** | **V** | **L** | **K** | **l-*allo*-Ile** |
| **Aba1 A8** | **D** | **A** | **W** | **M** | **L** | **L** | **A** | **V** | **I** | **K** | **l-Leu** |
| **GliP A2** | **D** | **Y** | **N** | **S** | **V** | **A** | **A** | **S** | **I** | **K** | **l-Ser** |
| **TqaA A3** | **D** | **M** | **V** | **I** | **I** | **L** | **G** | **S** | **A** | **K** | **l-Ala** |
| **Alb01 A1** | **S** | **V** | **K** | **Y** | **V** | **T** | **N** | **N** | **D** | **K** | **pABA** |
| **Alb01 A3** | **A** | **V** | **K** | **Y** | **V** | **T** | **N** | **N** | **D** | **K** | **pABA** |
| **NanA A2** | **G** | **A** | **G** | **M** | **L** | **L** | **G** | **T** | **V** | **K** | **l-Ky** |
| **RifA A1** | **D** | **L** | **I** | **A** | **G** | **A** | **A** | **G** | **A** | **K** | **AHBA** |
| **Aba1 A1** | **D** | **A** | **L** | **L** | **V** | **L** | **I** | **T** | **V** | **K** | **d-Hmp** |
| **TqaB A** | **D** | **L** | **F** | **M** | **V** | **L** | **G** | **G** | **C** | **K** | **AIA** |

**Table S4. Adenylation domain amino acid sequences analyzed in Figure S7.**

| **Protein**  **(domains)** | **Enzyme class** | **Function** | **Origin** | **Accession No.** |
| --- | --- | --- | --- | --- |
| PlmA  (A) | NRPS | Penilumamide synthetase I | *Aspergillus flavipes* CNL-338  (this work) | ON297638 |
| PlmJ  (A1,A2) | NRPS | Penilumamide synthetase II | *Aspergillus flavipes* CNL-338  (this work) | ON297638 |
| PlmK  (A) | NRPS | Penilumamide synthetase III | *Aspergillus flavipes* CNL-338  (this work) | ON297638 |
| Aba1  (A1-9) | NRPS | Aureobasidin A1 complex | *Aureobasidium pullulans* | ACJ04424.1 |
| Alb01  (A1,A3) | PKS-NRPS | Albicidin synthetase 1 | *Xanthomonas albilineans* | CAE52339.1 |
| AnaPS  (A1,A2) | NRPS | Acetylaszonalenin synthetase | *Aspergillus fischeri* NRRL 181 | A1DN09.1 |
| AuaEII  (A) | NRPS | anthranilate-CoA ligase | *Stigmatella aurantiaca Sg a15* | CCA65703.1 |
| BenY  (A) | NRPS | Benzomalvin synthetase Y | *Aspergillus terreus* | P9WEU8.1 |
| BenZ  (A1,A2) | NRPS | Benzomalvin synthetase Z | *Aspergillus terreus* | P9WEU9.1 |
| CcsA  (A) | PKS-NRPS | Cytochalasin synthetase A | *Aspergillus clavatus* NRRL 1 | A1CLY8 |
| CheA  (A) | PKS-NRPS | Chaetoglobosin synthetase A | *Penicillium expansum* | CAO91861.1 |
| FkbP  (A) | NRPS | FK506-binding protein | *Streptomyces hygroscopicus* subsp. *ascomyceticus* | AAF86395.1 |
| GetE  (A) | NRPS | GE81112 adenylation domain | *Streptomyces* sp. L-49973 | CBL93716.1 |
| GliP  (A1,A2) | NRPS | Gliotoxin synthetase | *Aspergillus fumigatus* | AAW03307.1 |
| GrsA  (A1) | NRPS | Gramicidin S synthetase A | *Brevibacillus brevis* | P0C062.1 |
| IvoA  (A) | NRPS | Ivory mutation-related protein A | *Aspergillus nidulans* FGSC A4 | C8V7P4.1 |
| NanA  (A1,A2) | NRPS | Nanangelenin synthetase A | *Aspergillus nanangensis* | QIQ51365.1 |
| NpsP8 | NRPS | Napsamycin biosynthesis protein | *Streptomyces* sp. DSM 5940 | ADY76684.1 |
| PsyA  (A1,A2) | NRPS | Psychrophilin synthetase A | *Penicillium* sp. YT-2016 | AMQ36132.1 |
| PsyC  (A) | NRPS | Psychrophilin synthetase C | *Penicillium* sp. YT-2016 | AMQ36134.1 |
| RapP  (A) | NRPS | Pipecolate incorporating enzyme | *Streptomyces hygroscopicus* | Q54298.1 |
| RifA  (A1) | PKS-NRPS | Rifamycin polyketide synthase A | *Amycolatopsis mediterranei* | O54666 |
| SnbA  (A) | NRPS | Pristinamycin I synthetase 1 | *Streptomyces pristinaespiralis* | P95819 |
| SnbDE  (A3) | NRPS | Pristinamycin I synthase 3 and 4 | *Streptomyces pristinaespiralis* | O07944 |
| TqaA  (A1,A2,A3) | NRPS | Tryptoquialanine synthetase A | *Penicillium aethiopicum* | ADY16697.1 |
| TqaB  (A) | NRPS | Tryptoquialanine synthetase B | *Penicillium aethiopicum* | ADY16689.1 |

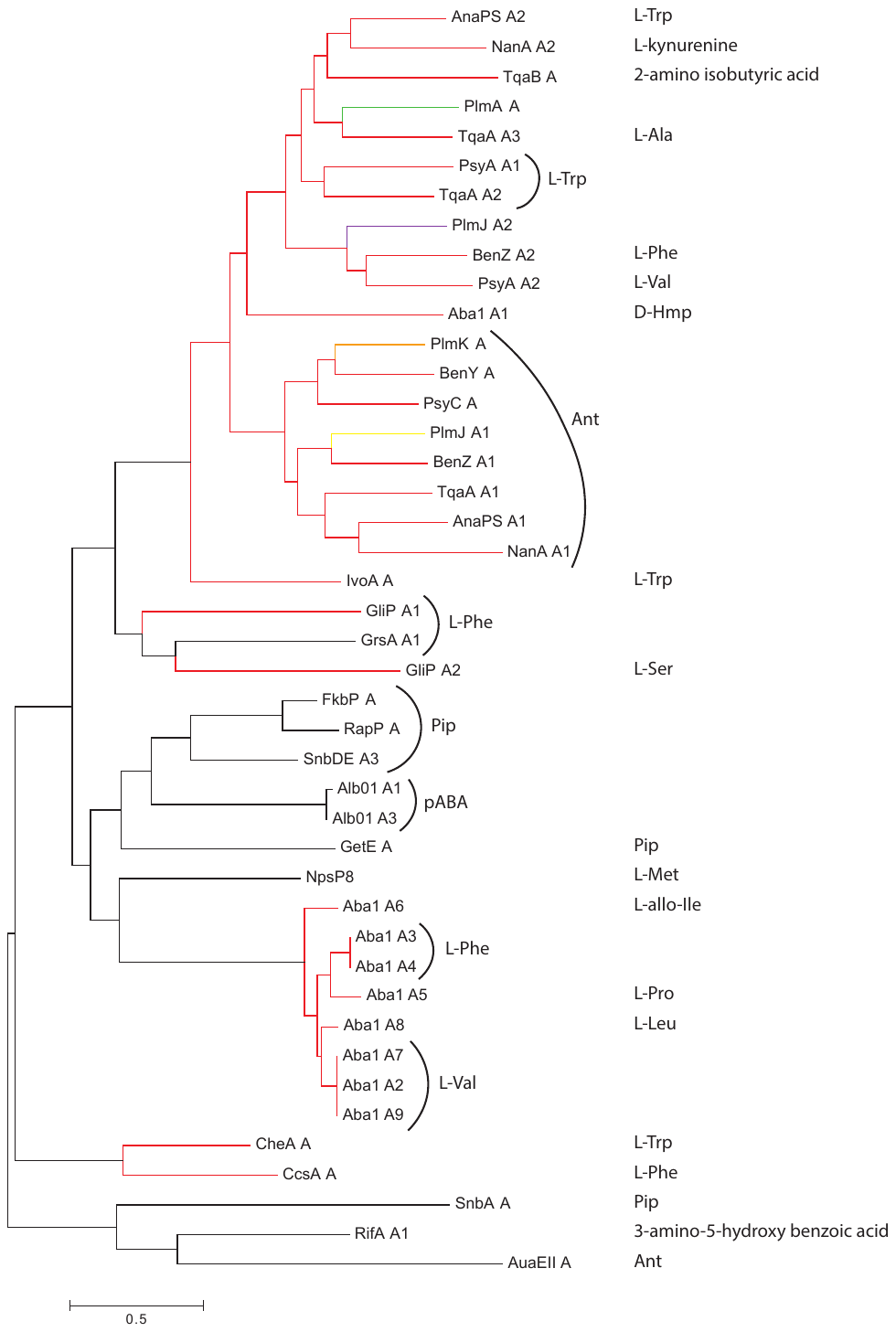

**Figure S7. Maximum-likelihood phylogenetic tree of 43 NRPS A domains based on substrate selectivity.** Black lines are bacterial A domains, and red lines are fungal A domains. The four *plm* A domains are in green (PlmA A), yellow (PlmJ A1), purple (PlmJ A2), and orange (PlmK A). The maximum-likelihood tree was generated using MEGA 6.0 software with the JTT model of amino acid substitution. The scale bar represents the average number of amino acid substitutions. d -Hmp = d-2-hydroxy-3-methylpentanoic acid, Ant = anthranilic acid, Pip = pipecolate, pABA = para-aminobenzoic acid.

**Table S5. Protein sequences used for the condensation domain comparison in Figure S8.**

| **Protein**  **(domains)** | **Enzyme class** | **Function** | **Origin** | **Accession No.** |
| --- | --- | --- | --- | --- |
| PlmA  (C1,C2) | NRPS | Penilumamide synthetase I | *Aspergillus flavipes* CNL-338  (this work) | ON297683 |
| PlmJ  (C1,C2) | NRPS | Penilumamide synthetase II | *Aspergillus flavipes* CNL-338  (this work) | ON297683 |
| PlmK  (C) | NRPS | Penilumamide synthetase III | *Aspergillus flavipes* CNL-338  (this work) | ON297683 |
| Aba1  (C1-9) | NRPS | Aureobasidin A1 complex | *Aureobasidium pullulans* | ACJ04424.1 |
| Alb01  (C1,C2) | PKS-NRPS | Albicidin synthetase 1 | *Xanthomonas albilineans* | CAE52339.1 |
| BenY  (C) | NRPS | Benzomalvin synthetase Y | *Aspergillus terreus* | P9WEU8.1 |
| BenZ  (C1,C2) | NRPS | Benzomalvin synthetase Z | *Aspergillus terreus* | P9WEU9.1 |
| FkbP  (C1,C2) | NRPS | FK506-binding protein | *Streptomyces hygroscopicus* subsp. *ascomyceticus* | AAF86395.1 |
| Fum14p  (C) | NRPS | Fumonisin biosynthesis protein 14 | *Fusarium verticillioides* | AAN74817.2 |
| GliP  (C1,C2) | NRPS | Gliotoxin synthetase | *Aspergillus fumigatus* | AAW03307.1 |
| GrsA  (C) | NRPS | Gramicidin S synthetase A | *Brevibacillus brevis* | P0C062.1 |
| GrsB  (C1-3) | NRPS | Gramicidin S synthetase B | *Brevibacillus brevis* | CAA434838.1 |
| IvoA  (C1,C2) | NRPS | Ivory mutation-related protein A | *Aspergillus nidulans* FGSC A4 | C8V7P4.1 |
| LNKS  (C) | PKS-NRPS | Lovastatin nonaketide synthase (LovB) | *Aspergillus terreus* | Q9Y8A5.1 |
| NanA  (C1,C2) | NRPS | Nanangelenin synthetase A | *Aspergillus nanangensis* | QIQ51365.1 |
| NpsP2 | NRPS | Napsamycin biosynthesis protein | *Streptomyces* sp. DSM 5940 | ADY76661.1 |
| NpsP4 | NRPS | Napsamycin biosynthesis protein | *Streptomyces* sp. DSM 5940 | ADY76666.1 |
| PsyA  (C1,C2) | NRPS | Psychrophilin synthetase A | *Penicillium* sp. YT-2016 | AMQ36132.1 |
| PsyC  (C) | NRPS | Psychrophilin synthetase C | *Penicillium* sp. YT-2016 | AMQ36134.1 |
| RapP  (C1,C2) | NRPS | Pipecolate incorporating enzyme | *Streptomyces hygroscopicus* | Q54298.1 |
| SbtI2  (C) | NRPS | Serobactin synthetase 2 | *Herbaspirillum seropedicae* | QDD64765.1 |
| SrfAA  (C1) | NRPS | Surfactin synthetase A | *Bacillus subtilis* subsp. *subtilis* str. 168 | NP_388230.2 |
| SyfB  (C3,C5) | NRPS | Syringafactin synthetase B | *Pseudomonas syringae* group *genomosp.* 3 | WP_011104220.1 |
| Tcp12  (C2) | NRPS | Teicoplanin synthetase | *Actinoplanes teichomyceticus* | Q70AZ6 |
| TqaA  (C1-4) | NRPS | Tryptoquialanine synthetase A | *Penicillium aethiopicum* | ADY16697.1 |
| TqaB  (C) | NRPS | Tryptoquialanine synthetase B | *Penicillium aethiopicum* | ADY16689.1 |
| TwmB  (C) | PKS-NRPS | Wortmanamide synthetase B | *Talaromyces wortmannii* | QBC19710.1 |
| VibF  (C1) | NRPS | Vibriobactin synthetase F | *Vibrio cholerae* | WP_001923521.1 |
| VibH | NRPS | Vibriobactin synthetase H | *Vibrio cholerae* | AAD48879.1 |

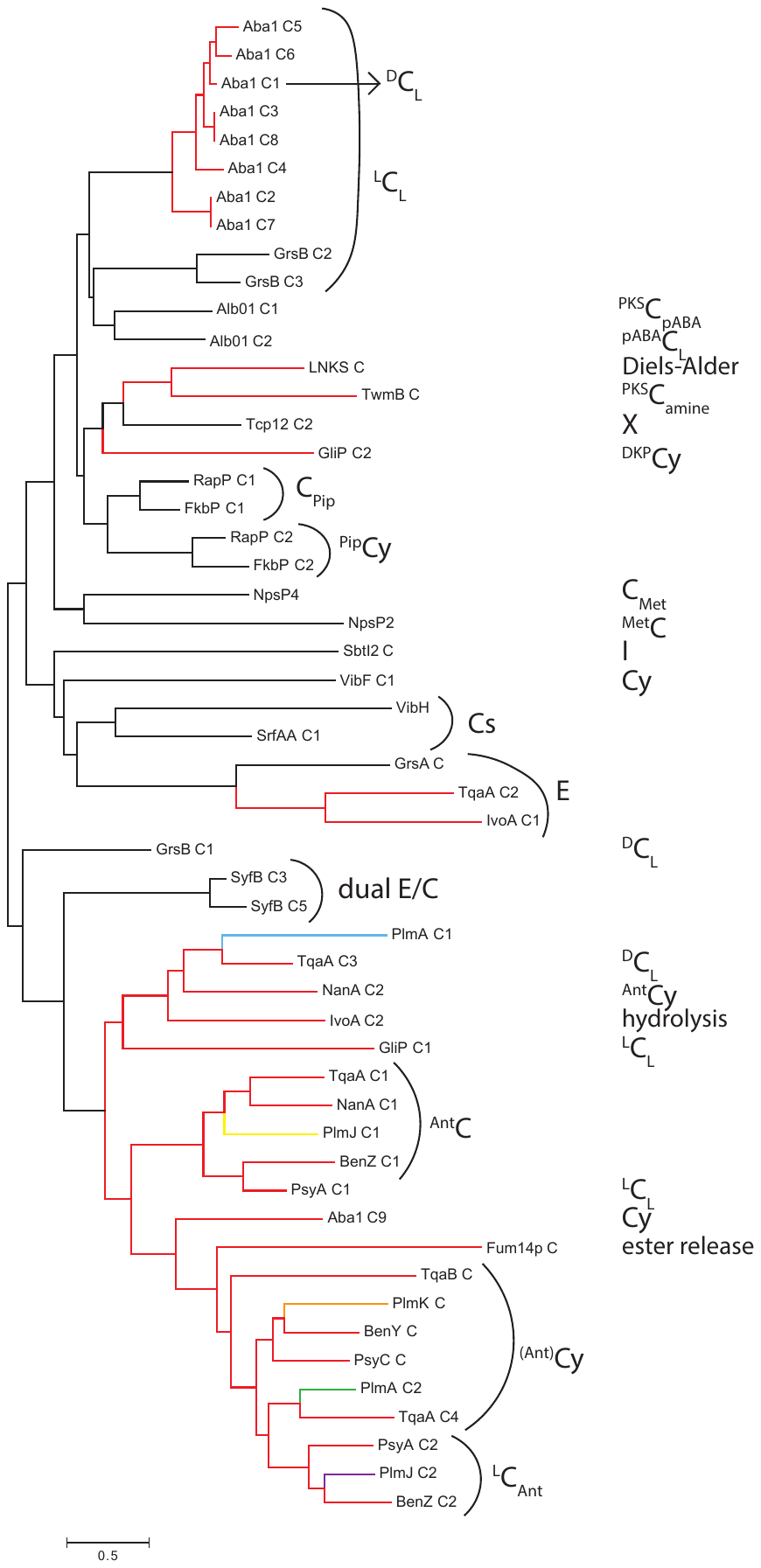

**Figure S8. Maximum-likelihood phylogenetic tree of 53 NRPS C domains based on function.** Black lines are bacterial C domains, and red lines are fungal C domains. The five *plm* C domains are in blue (PlmA C1), green (PlmA C2), yellow (PlmJ C1), purple (PlmJ C2), and orange (PlmK C). The maximum-likelihood tree was generated using MEGA 6.0 software with the JTT model of amino acid substitution. The scale bar represents the average number of amino acid substitutions. Cs = starter C domain, Cy = terminal cyclizing C domain, E = epimerization domain, I = interfacing domain, X = interfacing domain that recruits Oxy enzymes, and dual E/C = bifunctional C domain that also epimerizes the donor substrate. ^X^C_Y_ notation denotes the donor substrate as X and the acceptor substrate as Y. l/d = proteinogenic amino acid stereochemistry, PKS = polyketide synthase, pABA = para-aminobenzoic acid, DKP = diketopiperazine, Pip = pipecolate, Met = methionine, Ant = anthranilic acid.

**Table S6. Protein sequences used for the condensation domain comparison in Figure S9.**

| **Protein (domains)** | **Enzyme class** | **Function** | **Origin** | **Accession No.** |
| --- | --- | --- | --- | --- |
| PlmA  (C1,C2) | NRPS | Penilumamide synthetase I | *Aspergillus flavipes* CNL-338 | ON297683 |
| PlmJ  (C1,C2) | NRPS | Penilumamide synthetase II | *Aspergillus flavipes* CNL-338 | ON297683 |
| PlmK  (C) | NRPS | Penilumamide synthetase III | *Aspergillus flavipes* CNL-338 | ON297683 |
| Aba1  (C9) | NRPS | Aureobasidin A1 complex | *Aureobasidium pullulans* | ACJ04424.1 |
| AcmB  (C1,C2,C4) | NRPS | Actinomycin synthetase II | *Streptomyces anulatus* | O68487 |
| AcmC  (C1-3) | NRPS | Actinomycin synthetase III | *Streptomyces anulatus* | Q9L8H4 |
| AebF  (C) | NRPS | Enterobactin synthetase F | *Vibrio campbellii* | KGR33264.1 |
| AltG  (C) | NRPS | Bromoalterochromide interfacing domain | *Pseudoalteromonas luteoviolacea* | WP_063365585.1 |
| AltI  (C) | NRPS | Bromoalterochromide interfacing domain | *Pseudoalteromonas luteoviolacea* | WP_063365570.1 |
| AmbE  (C) | NRPS | AMB synthetase E | *Pseudomonas aeruginosa* PAK | VUY44935.1 |
| ArfA  (C1,C2) | NRPS | Arthrofactin synthetase A | *Pseudomonas* sp. MIS38 | BAC67534.2 |
| ArfC  (C2,C4) | NRPS | Arthrofactin synthetase C | *Pseudomonas* sp. MIS38 | Q84BQ4 |
| Bamb_5915  (C) | NRPS | Enacyloxin synthetase | *Burkholderia ambifaria* AMMD | ABI91460.1 |
| BpsA  (C1-3) | NRPS | Balhimycin synthetase A | *Amycolatopsis balhimycina* | Q939Z1 |
| BpsB  (C1-4) | NRPS | Balhimycin synthetase B | *Amycolatopsis balhimycina* | Q939Z0 |
| BpsC  (C1,C2) | NRPS | Balhimycin synthetase C | *Amycolatopsis balhimycina* | Q939Y9 |
| CdaPS1  (C1-4,C7) | NRPS | CDA synthetase 1 | *Streptomyces coelicolor* | WP_011028842.1 |
| CdaPS2  (C1-4) | NRPS | CDA synthetase 2 | *Streptomyces coelicolor* | WP_011028843.1 |
| CroK  (C2) | NRPS | Crochelin synthetase | *Chondromyces crocatus* | AIR74925.1 |
| CrpD  (C2) | NRPS | Cryptophycin synthetase D | *Nostoc* sp. ATCC 53789 | QHG20896.1 |
| Dbv16  (C1,C2) | NRPS | Glycopeptide synthetase  X domain | *Nonomuraea gerenzanensis* | Q7WZ75 |
| Dbv17  (C1-4) | NRPS | Glycopeptide synthetase  E domain | *Nonomuraea gerenzanensis* | Q7WZ74 |
| Dbv25  (C1,C2) | NRPS | Glycopeptide synthetase  E domain | *Nonomuraea gerenzanensis* | Q7WZ66 |
| Dbv26  (C) | NRPS | Glycopeptide synthetase | *Nonomuraea gerenzanensis* | Q7WZ65 |
| DhbF  (C1) | NRPS | Bacillibactin synthetase F | *Bacillus cereus* | WP_011198679.1 |
| EntF  (C) | NRPS | Enterobactin synthetase F | *Escherichia coli* | CAD6019783.1 |
| ErcD  (C5) | NRPS | Erythrochelin synthetase | *Saccharopolyspora erythraea* NRRL 2338 | CAM02313.1 |
| Fum14p  (C) | NRPS | Fumonisin biosynthesis protein 14 | *Fusarium verticillioides* | AAN74817.2 |
| GliP  (C2) | NRPS | Gliotoxin synthetase | *Aspergillus fumigatus* | AAW03307.1 |
| GrsA  (C) | NRPS | Gramicidin S synthetase A | *Brevibacillus brevis* | P0C062.1 |
| GrsB  (C1) | NRPS | Gramicidin S synthetase B | *Brevibacillus brevis* | P0C064.2 |
| GrsB  (C1-3) | NRPS | Gramicidin S synthetase B | *Brevibacillus brevis* | CAA434838.1 |
| HcsF  (C) | NRPS | Interfacing domain | *Pseudomonas thivervalensis* | WP_053122086.1 |
| HcsI  (C) | NRPS | Interfacing domain | *Pseudomonas thivervalensis* | WP_053122092.1 |
| HMWP2  (C1) | NRPS | Yersiniabactin synthetase | *Pseudomonas syringae* group *genomosp.* 3 | WP_011104107.1 |
| IcoA  (C3,C4) | NRPS | Icosalide synthetase | *Burkholderia gladioli* | AYA44686.1 |
| IvoA  (C2) | NRPS | Ivory mutation-related protein A | *Aspergillus nidulans* FGSC A4 | C8V7P4.1 |
| LicA  (C1,C4) | NRPS | Lichenysin synthetase A | *Bacillus licheniformis* | WP_011197536.1 |
| LicB  (C4) | NRPS | Lichenysin synthetase B | *Bacillus licheniformis* | WP_011197537.1 |
| LNKS  (C) | PKS-NRPS | Lovastatin nonaketide synthase (LovB) | *Aspergillus terreus* | Q9Y8A5.1 |
| MbtB  (C) | NRPS | Mycobactin synthetase B | *Mycobacterium avium* | WP_010949506.1 |
| MbtB  (C) | NRPS | Mycobactin synthetase B | *Mycobacterium tuberculosis* | WP_010950705.1 |
| McyA  (C) | NRPS | Microcystin synthetase A | *Microcystis aeruginosa* PCC 7806 | CAO90227.1 |
| MloJ  (C2) | PKS-NRPS | Malonomycin synthetase J | *Streptomyces rimosus* subsp. *paromomycinus* | AYJ71721.1 |
| NanA  (C2) | NRPS | Nanangelenin synthetase A | *Aspergillus nanangensis* | QIQ51365.1 |
| NbtF  (C) | NRPS | Mycobactin synthetase F | *Nocardia farcinica* | WP_011207299.1 |
| NocB  (C) | NRPS | Nocardicin synthetase B | *Nocardia uniformis* subsp. *tsuyamanensis* | AAT09805.1 |
| NosA  (C2,C3) | NRPS | Nostopeptolide  synthetase A | *Nostoc* sp. GSV224 | Q9RAH4 |
| NosC  (C1,C2) | NRPS | Nostopeptolide  synthetase C | *Nostoc* sp. GSV224 | Q9RAH2 |
| NosD  (C1,C2) | NRPS | Nostopeptolide  synthetase D | *Nostoc* sp. GSV224 | Q9RAH1 |
| NRPS  (C1,C2) | NRPS | Nonribosomal peptide synthetase | *Pectobacterium atrosepticum* | WP_011093071.1 |
| NRPS  (C1) | NRPS | Nonribosomal peptide synthetase | *Pseudomonas aeruginosa* PAO1 | NP_252017.1 |
| NRPS  (C1) | NRPS | Nonribosomal peptide synthetase | *Nocardia farcinica* | WP_011209327.1 |
| NRPS  (C1) | NRPS | Nonribosomal peptide synthetase | *Photobacterium profundum* | WP_011218392.1 |
| NRPS  (C1) | NRPS | Nonribosomal peptide synthetase | *Yersinia pseudotuberculosis* | WP_011193026.1 |
| NRPS  (C2) | NRPS | Nonribosomal peptide synthetase | *Yersinia pseudotuberculosis* | WP_011193027.1 |
| NRPS  (C2) | NRPS | Nonribosomal peptide synthetase | *Streptomyces coelicolor* | WP_011031827.1 |
| NRPS  (C1,C3) | NRPS | Nonribosomal peptide synthetase | *Chromobacterium violaceum* | WP_011136349.1 |
| NRPS  (C1,C3) | NRPS | Nonribosomal peptide synthetase | *Burkholderia pseudomallei* | WP_011205655.1 |
| NRPS  (C4) | NRPS | Nonribosomal peptide synthetase | *Pseudomonas protegens* | WP_011062464.1 |
| OfaA  (C1) | NRPS | Orfamide A synthetase A | *Pseudomonas protegens* | WP_011060446.1 |
| OfaB  (C2,C4) | NRPS | Orfamide A synthetase B | *Pseudomonas protegens* | WP_011060447.1 |
| OrbJ  (C2) | NRPS | Ornibactin synthetase J | *Burkholderia cepacia* | AUD11993.1 |
| PchE  (C) | NRPS | Pyochelin synthetase E | *Pseudomonas protegens* | ABW70809.1 |
| PchE  (C) | NRPS | Pyochelin synthetase E | *Pseudomonas aeruginosa* PAO1 | NP_252916.1 |
| PchF  (C1) | NRPS | Pyochelin synthetase F | *Burkholderia pseudomallei* | WP_011205430.1 |
| PchF  (C1) | NRPS | Pyochelin synthetase F | *Pseudomonas protegens* | WP_011061772.1 |
| PCZA363.3  (C1-3) | NRPS | Vancomycin group synthetase | *Amycolatopsis orientalis* | O52819 |
| PCZA363.4  (C1-4) | NRPS | Vancomycin group synthetase | *Amycolatopsis orientalis* | O52820 |
| PCZA363.5  (C1,C2) | NRPS | Vancomycin group synthetase | *Amycolatopsis orientalis* | O52821 |
| PfbI  (C) | NRPS | Interfacing domain | *Alcanivorax pacificus* | WP_052269209.1 |
| plu2320  (C1) | NRPS | Yersiniabactin synthetase | *Photorhabdus laumondii* | WP_011146562.1 |
| plu2670  (C6,C9) | NRPS | Nonribosomal peptide synthetase | *Photorhabdus laumondii* | WP_011146892.1 |
| plu2796  (C) | dehydrogenase-NRPS | Pepteridine synthetase | *Photorhabdus laumondii* subsp. *laumondii* TTO1 | CAE15170.1 |
| PpsA  (C1-3) | NRPS | Plipastatin synthetase A | *Bacillus subtilis* subsp. *subtilis* str. 168 | NP_389716.2 |
| PpsB  (C2,C3) | NRPS | Plipastatin synthetase B | *Bacillus subtilis* subsp. *subtilis* str. 168 | NP_389715.1 |
| PpsC  (C1-3) | NRPS | Plipastatin synthetase C | *Bacillus subtilis* subsp. *subtilis* str. 168 | NP_389714.1 |
| PpsD  (C2,C4) | NRPS | Plipastatin synthetase D | *Bacillus subtilis* subsp. *subtilis* str. 168 | NP_389713.2 |
| PpsE  (C) | NRPS | Plipastatin synthetase E | *Bacillus subtilis* subsp. *subtilis* str. 168 | NP_389712.1 |
| PSEEN_RS  14935 (C) | NRPS | Interfacing domain | *Pseudomonas entomophila* | WP_011534377.1 |
| PS1  (C1,C2) | NRPS | Peptide synthetase | *Streptomyces lavendulae* | Q93N86 |
| PS2  (C1-3,C6) | NRPS | Peptide synthetase | *Streptomyces lavendulae* | Q93N87 |
| PS3  (C1,C2) | NRPS | Peptide synthetase | *Streptomyces lavendulae* | Q93N88 |
| PS4  (C1,C2) | NRPS | Peptide synthetase | *Streptomyces lavendulae* | Q93N89 |
| Pvd2  (C2) | NRPS | Pyoverdine synthetase 2 | *Pseudomonas syringae* group *genomosp.* 3 | WP_011103871.1 |
| Pvd4  (C2) | NRPS | Pyoverdine synthetase 4 | *Pseudomonas syringae* group *genomosp.* 3 | WP_011103873.1 |
| PvdD  (C3) | NRPS | Pyoverdine synthetase D | *Pseudomonas protegens* | WP_011062374.1 |
| PvdI  (C4) | NRPS | Pyoverdine synthetase I | *Pseudomonas protegens* | WP_011062376.1 |
| PvdJ  (C) | NRPS | Pyoverdine synthetase J | *Pseudomonas taiwanensis* | AJW67534.1 |
| PvdK  (C) | NRPS | Pyoverdine synthetase K | *Pseudomonas fluorescens* | WP_011333311.1 |
| RapP  (C1,C2) | NRPS | Pipecolate incorporating enzyme | *Streptomyces rapamycinicus* NRRL 5491 | BCH36730.1 |
| SbtI2  (C) | NRPS | Serobactin synthetase 2 | *Herbaspirillum seropedicae* | QDD64765.1 |
| SbtI2  (C) | NRPS | Serobactin synthetase 2 | *Herbaspirillum seropedicae* | WP_048348543.1 |
| SfmC  (C) | NRPS | Saframycin synthetase C | *Streptomyces lavendulae* | ABI22133.1 |
| SgcC5 | NRPS | C-1027 condensation domain | *Streptomyces* sp. CB02366 | ANY94448.1 |
| SnbC  (C1-3) | NRPS | Pristinamycin I synthetase 2 | *Streptomyces pristinaespiralis* | Q54959 |
| SnbDE  (C1-3) | NRPS | Pristinamycin I synthase  3 and 4 | *Streptomyces pristinaespiralis* | O07944 |
| SrfAA  (C1,C2,C4) | NRPS | Surfactin synthetase A | *Bacillus subtilis* subsp. *subtilis* str. 168 | NP_388230.2 |
| SrfAB  (C1,C2,C4) | NRPS | Surfactin synthetase B | *Bacillus subtilis* subsp. *subtilis* str. 168 | NP_388231.2 |
| SrfAC  (C) | NRPS | Surfactin synthetase C | *Bacillus subtilis* subsp. *subtilis* str. 168 | NP_388233.2 |
| StaA  (C1,C2) | NRPS | Staurosporine synthetase A | *Streptomyces toyocaensis* | Q8KLL3 |
| StaB  (C1,C2) | NRPS | Staurosporine synthetase B | *Streptomyces toyocaensis* | Q8KLL4 |
| StaC  (C1-4) | NRPS | Staurosporine synthetase C | *Streptomyces toyocaensis* | Q8KLL5 |
| StaD  (C1,C2) | NRPS | Staurosporine synthetase D | *Streptomyces toyocaensis* | Q8KLL6 |
| SyfA  (C1,C2) | NRPS | Syringafactin synthetase A | *Pseudomonas syringae* group *genomosp.* 3 | WP_011104219.1 |
| SyfB  (C3,C5) | NRPS | Syringafactin synthetase B | *Pseudomonas syringae* group *genomosp.* 3 | WP_011104220.1 |
| SyrE  (C5,C6) | NRPS | Syringomycin synthetase | *Pseudomonas syringae* pv. *syringae* | O85168 |
| TaiE  (C) | PKS-NRPS | Thailandamide  synthetase E | *Cupriavidus taiwanensis* | WP_012356046.1 |
| Tcp9  (C1,C2) | NRPS | Teicoplanin synthetase | *Actinoplanes teichomyceticus* | Q70AZ9 |
| Tcp10  (C) | NRPS | Teicoplanin synthetase | *Actinoplanes teichomyceticus* | Q70AZ8 |
| Tcp11  (C1-4) | NRPS | Teicoplanin synthetase | *Actinoplanes teichomyceticus* | Q70AZ7 |
| Tcp12  (C1,C2) | NRPS | Teicoplanin synthetase | *Actinoplanes teichomyceticus* | Q70AZ6 |
| Tcp12  (C2) | NRPS | Teicoplanin synthetase | *Actinoplanes teichomyceticus* | CAE53353.1 |
| TqaA  (C4) | NRPS | Tryptoquialanine  synthetase A | *Penicillium aethiopicum* | ADY16697.1 |
| TwmB  (C) | PKS-NRPS | Wortmanamide  synthetase B | *Talaromyces wortmannii* | QBC19710.1 |
| TycB  (C1) | NRPS | Tyrocidine synthetase B | *Brevibacillus brevis* | QDS34188.1 |
| Var5  (C) | NRPS | Interfacing domain | *Variovorax paradoxus* | ALG65340.1 |
| VarH  (C) | NRPS | Interfacing domain | *Variovorax boronicumulans* | WP_062469880.1 |
| VibF  (C1) | NRPS | Vibriobactin synthetase F | *Vibrio cholerae* | WP_001923521.1 |
| VibH | NRPS | Vibriobactin synthetase H | *Vibrio cholerae* | AAD48879.1 |
| Zmn19 | NRPS | Zeamine condensation domain | *Serratia plymuthica* RVH1 | CCM44339.1 |

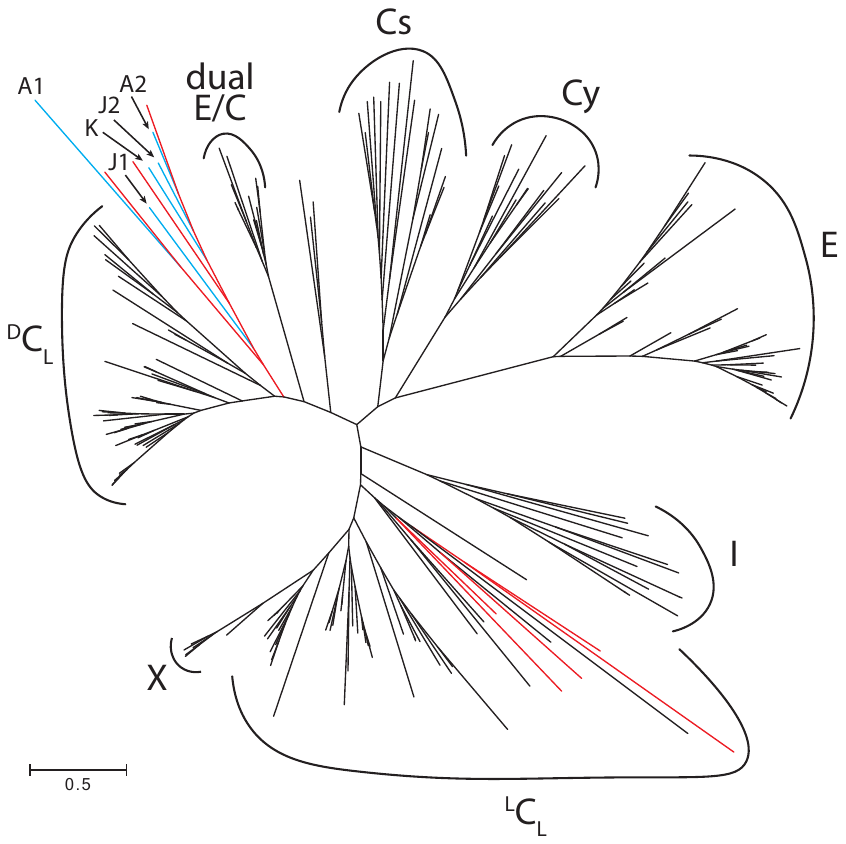

**Figure S9. Maximum-likelihood phylogenetic tree of 200 NRPS C domains based on function.** Modified from refs. 14 and 15. Black lines are bacterial C domains, red lines are fungal C domains, and blue lines are the five *plm* C domains: PlmA C1 (A1), PlmA C2 (A2), PlmJ C1 (J1), PlmJ C2 (J2), and PlmK C (K). The phylogenetic tree was generated using MEGA 6.0 software with the JTT model of amino acid substitution. The scale bar represents the average number of amino acid substitutions. Cs = starter C domain, Cy = terminal cyclizing C domain, E = epimerization domain, I = interfacing domain, ^L^C_L_ = C domain that condenses two l-amino acid substrates, X = interfacing domain that recruits P450 enzymes, ^D^C_L_ = C domain that condenses a d-amino acid donor with an l-amino acid acceptor, dual E/C = bifunctional C domain that also epimerizes the donor substrate.
